# Development of robust antiviral assays using relevant apical-out human airway organoids

**DOI:** 10.1101/2024.01.02.573939

**Authors:** Ji-Hoon Lee, Julia C. LeCher, Eric Parigoris, Noriyuki Shinagawa, Jason Sentosa, Candela Manfredi, Shu Ling Goh, Ramyani De, Sijia Tao, Keivan Zandi, Franck Amblard, Eric J. Sorscher, Jason R. Spence, Rabindra Tirouvanziam, Raymond F. Schinazi, Shuichi Takayama

## Abstract

While breakthroughs with organoids have emerged as next-generation *in vitro* tools, standardization for drug discovery remains a challenge. This work introduces human airway organoids with reversed biopolarity (AORBs), cultured and analyzed in a high-throughput, single-organoid-per-well format, enabling milestones towards standardization. AORBs exhibit a spatio-temporally stable apical-out morphology, facilitating high-yield direct intact-organoid virus infection. Single-cell RNA sequencing and immunohistochemistry confirm the physiologically relevant recapitulation of differentiated human airway epithelia. The cellular tropism of five severe acute respiratory syndrome coronavirus 2 (SARS-CoV-2) strains along with host response differences between Delta, Washington, and Omicron variants, as observed in transcriptomic profiles, also suggest clinical relevance. Dose-response analysis of three well-studied SARS-CoV-2 antiviral compounds (remdesivir, bemnifosbuvir, and nirmatrelvir) demonstrates that AORBs efficiently predict human efficacy, comparable to gold-standard air-liquid interface cultures, but with higher throughput (∼10-fold) and fewer cells (∼100-fold). This combination of throughput and relevance allows AORBs to robustly detect false negative results in efficacy, preventing irretrievable loss of promising lead compounds. While this work leverages the SARS-CoV-2 study as a proof-of-concept application, the standardization capacity of AORB holds broader implications in line with regulatory efforts to push alternatives to animal studies.

## Introduction

Organoids recapitulate physiological tissue functions and reflect human biological variability, such as responsiveness to different therapeutics^1^, thus playing a critical role in the drug development pipeline. The functional features of organoids arise from stem or progenitor cell differentiation and assembly within a basement membrane protein-rich extracellular matrix (ECM) milieu. Organoid design typically includes a large excess of ECM, resulting in undesirable variability in cellular self-organization and leading to organoids of various shapes, sizes, and numbers (**Supplementary Fig. 1**). In lung organoid virus infection studies, the excess ECM surrounding the organoids and their generally apical-in structure limit the access of viruses and therapeutics to apically expressed receptors (**Supplementary Fig. 1e**)^2–5^. Furthermore, organoid formation tends to result in a solid cellular core, leading to hypoxia and necrosis, thereby limiting their lifespan. Therefore, despite their proven physiological tissue functionality, these limitations can hinder the scalability, standardization, and broader use of lung organoids in studies of the normal and diseased human airway and therapeutic development.

This paper describes a bioengineering solution to these challenges based on organoid self-organization in minimal ECM to generate stable, apical-out human airway organoids in a one-organoid, one-well format (**Fig. 1a**). Unlike conventional organoid cultures, where macroscopic ECM domes encase the organoid exterior (**Fig. 1b**)^6^, our method results in cells surrounding a minimal amount of Matrigel scaffolding compartmentalized within the organoid interior^7–10^. While apical-out lung organoids can be produced from conventional cultures through the removal of extra-organoid ECM to trigger eversion^11,12^, the structures are small, often incompletely everted and appear inherently less stable, perhaps due to the removal of ECM. Importantly, direct organoid viral infection rates reported in the literature are inconsistent, or even often omitted^13^. This limitation stems from a fundamental challenge in conventional Matrigel dome culture: difficulty in controlling the number of organoids within each well and ensuring consistent viral delivery to the apical surface (**Supplementary Fig. 1**). As a result, accurately quantifying viral infection rates becomes a challenging task. In contrast, our ECM-encapsulating organoids are produced in a one-organoid, one-per-well manner using an eversion-free mechanism (**Supplementary Fig. 2**). They not only possess a clearly defined basal-in, stable apical-out structure that successfully remains structurally sustainable for over 60 days (**Supplementary Fig. 3**), but also are robustly and reproducibly cultured across five biological donors (**Table 1**). In addition, our organoids can be scaled up to 384-well plate for use in high-throughput screening (HTS) platforms. For brevity, we refer to these three-dimensional (3D) cellular constructs as “airway organoids with reversed biopolarity” (AORBs).

**Fig. 1.**
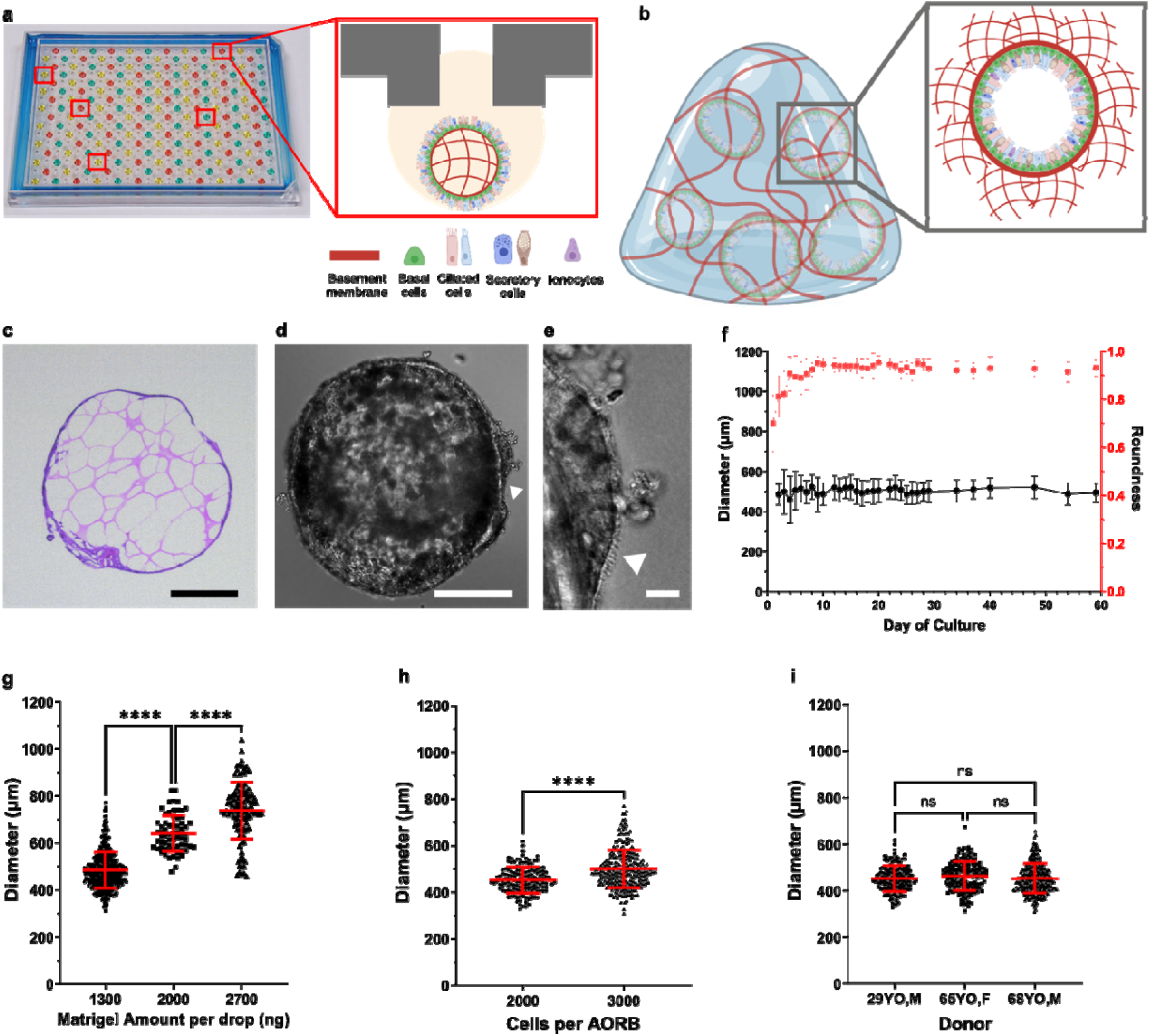
Human airway epithelial organoid with reversed biopolarity (AORBs). **a**, A 192 hanging drop array culture of single AORB-per-drop provides throughput and consistency. Eversion-free stably-inverted AORBs have terminally differentiated and physiologically relevant cell types. **b**, Conventional Matrigel-dome method of organoid culture has an exterior basement membrane with the apical side facing the interior. **c**, H&E staining shows an epithelial layer shell with a core filled with Matrigel. **d**, Phase-contrast image of an entire AORB, and **e**, a zoomed-in image showing cilia (denoted by triangles) and AORB-produced mucus. **f**, The diameters of AORBs are stable over 80 days, whereas the roundness increases up to day 10 of culture then stabilizes, *n* = 16 AORBs per timepoint. **g**, For a given number of starting cells, Matrigel amount can alter AORB diameters, *n* = 434, 66, 116 individual AORBs for 1300, 2000, and 2700 ng, respectively. **h**, Seeding density also provides a statistically significant means of regulating AORB diameters, *n* = 147 and 287 AORBs for 2000 and 3000 seeding cells, respectively. **i**, For a given Matrigel amount and cell density, AORB diameters are consistent across biological donors, *n* = 147, 157, and 290 for the three donors, respectively. ns, p>0.05, ****p<0.0001. Error bars indicate the mean ± 1 standard deviation. Scale bars, 250 µm in **c** and **d**, and 20 µm in **e**.

**Table 1.** Donor information for hBTECs from Lonza used in this study.

|  | Age | Sex | Race | Catalog#<br>(Lot#) |
| --- | --- | --- | --- | --- |
| Donor #1 | 29 | M | Caucasian | CC-2540S<br>(0000493462) |
| Donor #2 | 65 | F | Caucasian | CC-2540S<br>(0000619260) |
| Donor #3 | 68 | M | Caucasian | CC-2540S<br>(18TL134625) |
| Donor #4 | 21 | M | Caucasian | CC-2540S<br>(21TL244297) |
| Donor #5 | 73 | F | Caucasian | CC-2540S<br>(22TL290876) |

AORBs consist of physiologically relevant, *in vivo*-like, differentiated human respiratory epithelial cells, confirmed through single-cell RNA sequencing (scRNAseq). AORBs are readily infected with high yield (>95%) by five severe acute respiratory syndrome coronavirus 2 (SARS-CoV-2) strains. The stable apical-out design allows direct access of the virus to the apical face, while absence of extra-organoid ECM gel facilitates virus transport as well as exposure to antiviral compounds. Each AORB contains sufficient cells for single-organoid downstream quantitative reverse transcription polymerase chain reaction (RT-qPCR) analysis of virus infection. During evaluation of the antiviral activity of three anti-SARS-CoV-2 drugs, two of which are Food and Drug Administration (FDA)-approved and one in a phase III trial^14^, the antiviral efficacy results of AORBs reflect human efficacy even when conventional assays with 2D cell lines fail to show activity (**Table 2**). While the gold-standard transwell 24-well air-liquid interface (ALI) cultures also predict human efficacy, the AORB assays are significantly higher in throughput and at a 10x reduced cost. Moreover, screening proprietary compounds demonstrated the robust capability of AORBs to identify false negatives (missed hits) as well as false positives, validating their utility as a secondary screening platform.

**Table 2.** Anti-SARS-CoV-2 activities of four tested drugs in conventional 2D cell cultures, Transwell hBTEC-ALI culture, and AORBs.

| Compound | 2D Immortalized Culture Systems<sup>a)</sup> |  |  |  |  |  | 3D Primary Culture Systems<sup>a)</sup> |  |  |  |
| --- | --- | --- | --- | --- | --- | --- | --- | --- | --- | --- |
|  | Vero |  | Caco-2 |  | Calu-3 |  | hBTEC-ALI |  | AORBs |  |
|  | EC<sub>50</sub><br>(μM) | EC<sub>90</sub><br>(μM) | EC<sub>50</sub><br>(μM) | EC<sub>90</sub><br>(μM) | EC<sub>50</sub><br>(μM) | EC<sub>90</sub><br>(μM) | EC<sub>50</sub><br>(μM) | EC<sub>90</sub><br>(μM) | EC<sub>50</sub><br>(μM) | EC<sub>90</sub><br>(μM) |
| Remdesivir | 1.8 ± 0.1 | 2.6 ± 0.2 | 0.03 ± 0.01 | 0.08 ± 0.02 | 0.2 ± 0.1 | 0.4 ± 0.2 | 0.4 ± 0.3 | 0.7 ± 0.5 | 0.6 ± 1.0 | 2.3 ± 3.7 |
| AT-511 | 4.1 - >10 | >10 | >20 | >20 | 3.9 - >10 | >20 | 0.3 ± 0.3 | 0.5 ± 0.7 | 0.8 ± 1.0 | 1.8 ± 1.1 |
| Nirmatrelvir | 2.6 ± 0.6 | 4.2 ± 0.8 | 0.08 ± 0.004 | 0.015 ± 0.01 | 3.8 ± 1.1 | 11.1 ± 1.1 | 0.05 ± 0.003 | 0.2 ± 0.1 | 0.7 ± 1.4 | 1.3 ± 1.0 |

<sup>a)</sup>EC = 50% (EC50) or 90% (EC90) effective antiviral concentration. All values shown are mean ± standard deviation.

The field of everted- and inverted-organoids is a growing sub-field within the general organoid arena. Despite using similar starting cells, media, and supplements, the different material manipulations not only result in an inside-out geometry but also often lead to altered cellular differentiation. The demonstrated differentiation into all major bronchial epithelial cell types, stable inversion, and expression of the appropriate receptors and co-receptors for SARS-CoV-2 on the exterior, marks a significant advance in the manufacture of functional inverted airway organoids. Matrigel encapsulation by the epithelial cells stimulates the cellular production of a native basement membrane on the organoid interior, enhancing inversion stability. This ECM-encapsulating structure also allows for large-diameter organoids without a necrotic core, as most cells are on the surface, ensuring proper nutrient and oxygen diffusion. These features underscore the bioengineering significance of this work, enabling HTS in a one-organoid-per-well format (**Fig. 1a and Supplementary Fig. 2c**) for studies of the human airway and therapeutic development.

## Results

### Minimal Matrigel scaffolding supports high-throughput, robust, and single-organoid-per-well culture of AORBs

We previously described eversion-free methods for generating geometrically-inverted mammary organoids^7,8,10^. Additionally, we successfully adapted the methods for culturing kidney proximal tubule organoids, stably maintaining their inverted structure for over 100 days^9^. A key material manipulation method is to ’prematurely’ gel minimal Matrigel using warm media to culture the cells with sufficient, but not excessive, growth factors to promote cell assembly and epithelial sheet formation^8^. AORB production builds upon these general principles but was applied here to human bronchial/tracheal epithelial cells (hBTECs) cultured in 384-array hanging drop (HD) plates^15^ (**Supplementary Fig. 2**). Unique considerations for the use of hBTECs included identifying appropriate media conditions for initial AORB formation, as well as subsequent cellular differentiation. Unlike conventional apical-in organoids with a basement membrane that surrounds the organoid exterior (**Fig. 1b**), AORBs not only compartmentalize Matrigel in their core but also produce their own basement membrane on the interior surface (**Figs. 1c-e**). Optimization experiments culminated in a standard operating procedure (SOP) for AORB formation and culture (**Supplementary Fig. 3**).

The optimized SOP uses 3,000 cells per 25 µL droplet, Matrigel (50-107 μg/ml), fetal bovine serum (FBS) (<2%), and methylcellulose (0.24%) in a base medium. We achieved high-throughput (up to 384/plate; ∼210k organoids per donor) and robust AORB culture in a one-AORB-per-well format (**Supplementary Fig. 2**). AORBs remained stable for at least 60 days (**Supplementary Fig. 4**) with the diameter and roundness plateauing at consistent values (**Fig. 1f**). The compositions of the AORB seeding/formation and differentiation media are summarized in **Supplementary Table 1**. Analysis of >1,000 AORBs shows straightforward diameter modulation by adjusting Matrigel concentration (**Fig. 1g**) and the number of cells seeded (**Fig. 1h**). This observation held true across five donors (**Fig. 1i and Supplementary Figs. 5a-e; Table 1**) and across batches (**Supplementary Fig. 5f**), supporting the robustness of our SOP.

### AORBs comprise physiologically relevant differentiated airway cell types

Geometrical inversion was evidenced by exterior localization of apical markers acetylated α-tubulin (α-tub) on ciliated cells (**Fig. 2a and Supplementary Figs. 6a-e**) and mucin-5AC (MUC5AC) from goblet cells (**Fig. 2b**). In addition, live cell imaging confirmed beating apical-faced cilia (**Supplementary Videos 1-4**). Integrin beta-4 (ITGβ4) (**Fig. 2b**) and integrin alpha 6 (ITGα6) (**Fig. 2c**) were localized towards the interior of AORBs, confirming the inverted polarity. Images at lower magnification showed the stable global distribution of markers throughout AORBs (**Supplementary Figs. 6a-c and 7a-e**). We note that it is not uncommon for differentiated organoids to lose their basal progenitor cell population^16,17^, as they differentiate into different subtype airway cells (**Supplementary Figs. 8b**). In AORBs, however, keratin 5 (KRT5) staining showed sustained presence of correctly oriented basal cells together with the differentiated cells, reminiscent of physiological cell heterogeneity and tissue structure (**Fig. 2c**).

**Fig. 2.**
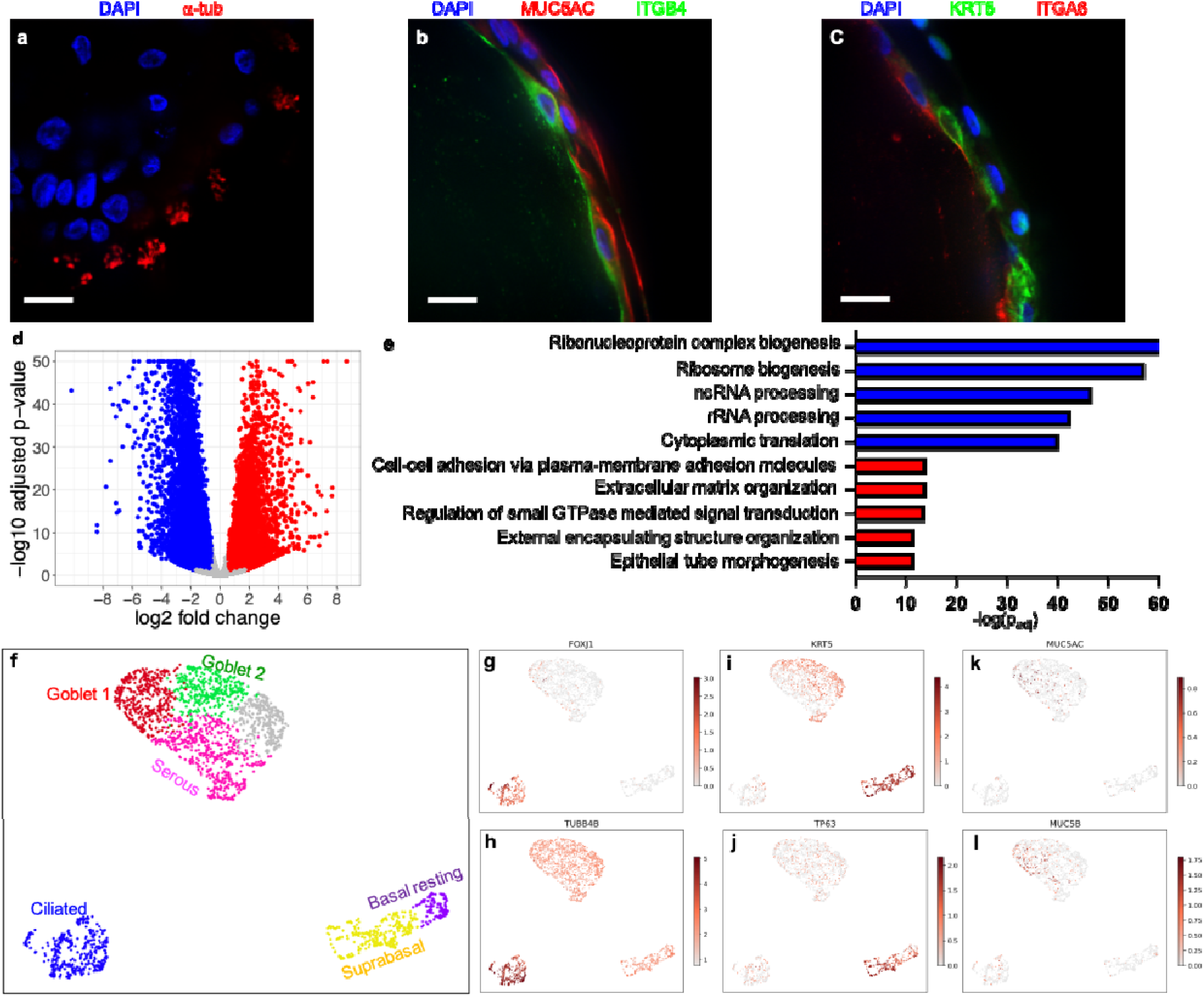
Confocal Immunofluorescence imaging confirms heterogeneous cell population and apical-out polarization in AORBs. **a**, Acetylated alpha tubulin orienting outwards indicating that the apical side of the ciliated cells face the organoid exterior. **b**, AORBs also have positive MUC5AC (red) indicative of goblet cells. ITGB4 (green) binds the basement membrane and indicates the basolateral side is facing the organoid interior. **c**, KRT5 (green) and ITGA6 (red) stains indicate the presence of basal cells. **d**, Volcano plot from bulk RNA sequencing comparing transcriptomic profile of the differentiated AORBs with respect to the initial cells before seeding (2D cultures) shows 7521 upregulated and 7100 downregulated genes. **e**, Top 5 statistically significant upregulated (red) and downregulated (blue) biological processes from gene ontology analysis of the enriched gene sets from DGE analysis. **f**, UMAP of scRNAseq results (*n* = 2651 cells) of AORBs showing three main clusters, ciliated, basal, and secretory. Further clustering shows sub-cell types, reprehensive of human bronchi. **g**-**l**, Canonical markers of ciliated cells (**g** and **h**), basal cells (**i** and **j**), and goblet cells (**k** and **l**). Color bars represent log-transformed normalized expression. Scale bars, 20 µm, in **a-c**.

Bulk RNAseq differential gene expression (DGE) analysis comparing AORB to their 2D undifferentiated counterparts revealed 7521 upregulated genes and 7100 downregulated genes (**Fig. 2d and Supplementary Fig. 8a; Supplementary Data 1**). AORBs showed downregulation of basal cell markers (**Supplementary Fig. 8b**), and upregulation of ciliated cell markers (**Supplementary Fig. 8c**). However, statistical tests suggested that the canonical markers of goblet cells (*MUC5AC*, *MUC5B*) were not differentially expressed in AORBs (**Supplementary Fig. 8d**); notably however, membrane-tethered mucins (*MUC1*, *MUC4*, *MUC16*) were upregulated (**Supplementary Fig. 8d**). Also notably, there was an upregulation of gene signatures of ionocytes (**Supplementary Fig. 8e**), a rare and understudied cell type of the human respiratory epithelium with unique expression of cystic fibrosis transmembrane conductance regulator protein^18,19^, further suggesting physiological relevance of our AORBs.

Gene ontology (GO) analysis was performed to assess enriched biological processes (**Fig. 2e**; **Supplementary Data 2**). Upregulated processes included cell-cell adhesions, ECM organization, GTPase-mediated signaling, and epithelial tube morphogenesis. Downregulated processes included rRNA processing, rRNA metabolism, and ribosome biogenesis, implying a differentiated state of AORBs^20^. The reduced proliferative state of AORBs was confirmed with negative log2FoldChange (LFC) of *MKI67*, *PCNA*, *MCM2*, and *CDK1* (**Supplementary Fig. 8f**). ECM organization-associated genes were also upregulated (**Supplemental Information**). Overall, DGE analysis implied the maturation and differentiation of AORBs. Results from signaling pathway impact analysis (SPIA) further substantiate that pathways for focal adhesion and ECM-receptor interaction were activated, whereas cell cycle and protein processing in the endoplasmic reticulum were inhibited (**Supplementary Data 3**).

scRNAseq of AORBs identified clusters of physiologically relevant cell types (**Figure 2f**). Representative canonical markers of ciliated (**Figures 2g-h**), basal (**Figures 2i-j**), and goblet cells (**Figures 2k-l**) were all enriched in the corresponding clusters. Reference-based mapping of the results based on the Human Lung Cell Atlas (HLCA)^21^ also showed relevant markers of cell types in these clusters (**Supplementary Figs. 8g-n**). Collectively, these data validated conversion of cultures from dividing basal epithelial progenitors to a fully differentiated, physiologically relevant representation of the human airway. Furthermore, the proportions of ciliated, basal, and goblet cells were approximately 20% for each. These proportions are consistent with those observed *in vivo*^22^, albeit slightly lower.

### IL-13 induces apical-out mucus hypersecretion

Interleukin-13 (IL-13) is a cytokine whose role in goblet cell metaplasia and mucus hypersecretion has been extensively studied^23–25^. To explore the functional aspect of the apical-out mucus secretion, AORBs were treated with 1 ng/mL IL-13 for 72 h (**Fig. 3a)** and mucins observed via bright field microscopy (**Fig. 3b**) and fluorescent lectin assays (**Fig. 3c and Supplementary Fig. 9**). Time-lapse imaging captured beating cilia clearing mucus on the apical surface (**Supplementary Videos 3-4**). In contrast, untreated AORBs showed qualitatively less mucus (**Fig. 3d**), suggesting stimulation of mucus hypersecretion by IL-13 (**Fig. 3e**). DGE analysis of IL-13-treated AORBs vs. untreated AORBs (**Supplementary Data 4)** confirmed upregulation of positive regulators of mucus secretion (*ANO1*, *ADORA3*, *ALOX12B*) and previously reported IL-13-induced mucus metaplasia related genes (*SPDEF*, *FOXA3*, *MUC2*, *MUC6*, *MUC5AC*, *CLCA1*, *ALOX15B*)^26^, and downregulation of negative regulators of mucus secretion (*ADA*) and ciliated cell markers (*TUBA1A*, *FOXJ1*) (**Fig. 3f**).

**Fig. 3.**
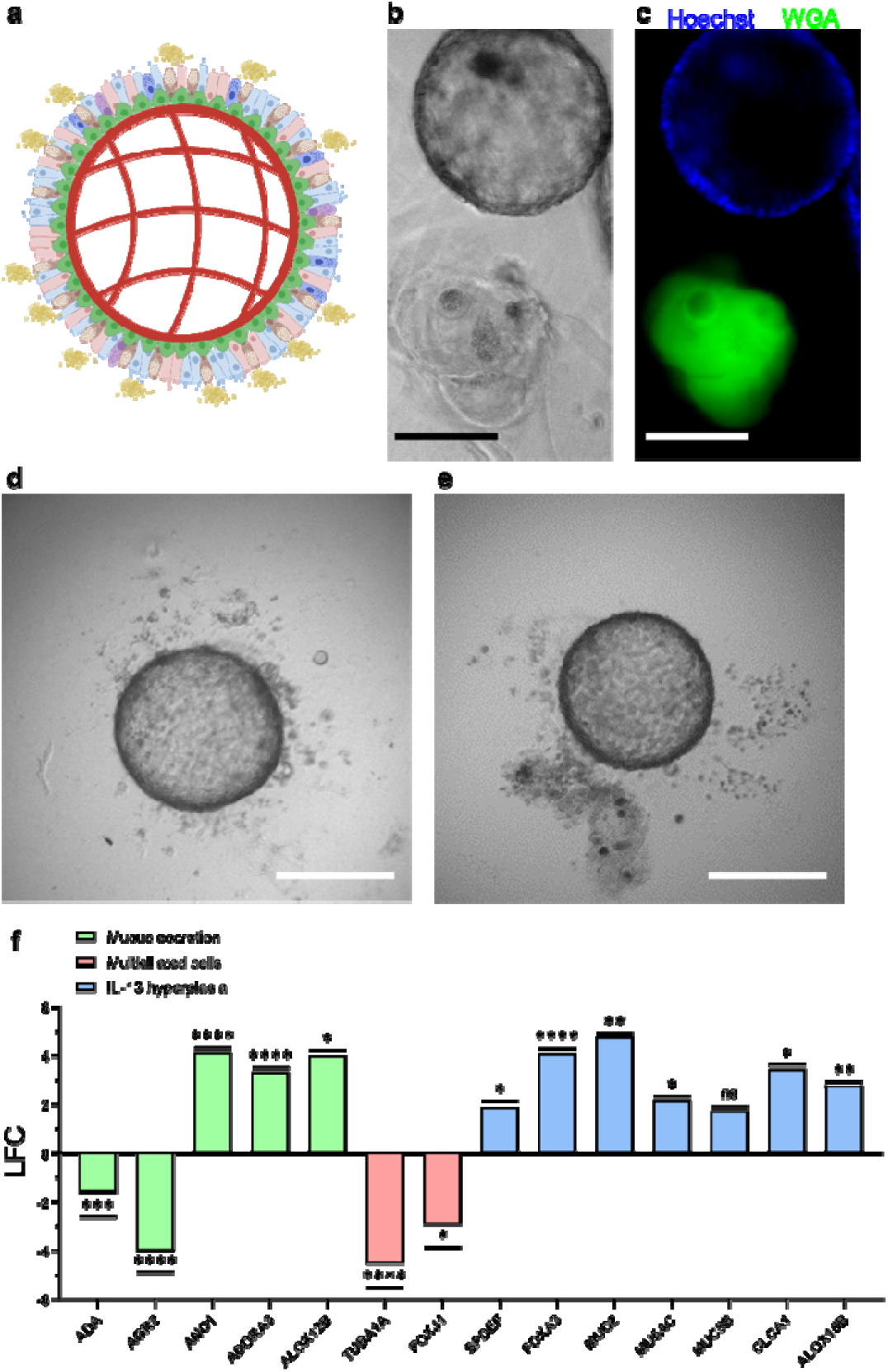
IL-13-induced apical-out mucus hypersecretion experiment. **a,** Apical-out morphology in AORBs allows efficient assay to study mucus secretion. **b** and **c**, Bright-field and immunofluorescent images of an AORB treated with 1 ng/mL IL-13, respectively. Nuclei and mucus are stained with Hoechst 33342 (blue) and wheat germ agglutinin (green). **d**, Representative bright-field images of untreated AORBs. **e**, Representative bright-field images of IL-13-treated. **f**, DGE analysis results (IL-13-treated vs. untreated) for genes associated with mucus secretion, multiciliated cells, and IL-13 hyperplasia, *n* = 3 biological donors per experimental condition. Scale bars, 500 µm in **b**-**e**.

### AORBs express SARS-CoV-2 entry-associated genes

It is well established that SARS-CoV-2 uses angiotensin-converting enzyme 2 (ACE2) and serine protease transmembrane serine protease 2 (TMPRSS2) for host cell entry. Immunofluorescence microscopy confirmed expression of ACE2 and TMPRSS2 on the apical surface of AORBs (**Fig. 4a** and **Supplementary Figs. 7f-g**). Complementary scRNAseq analysis confirmed expression of *ACE2* and *TMPRSS2* genes (**Figs. 4c-d**). ACE2 and TMPRRS2 expression steadily increased over 14 days, followed by a plateau (**Fig. 4b and Supplementary Fig. 7**). Moreover, TMPRSS2 had a diffuse pattern of staining (**Fig. 5b**), while ACE2 was apically-restricted (**Fig. 5c**), in agreement with previous observations that while TMPRSS2 is expressed throughout the membrane of airway epithelial cells (**Supplementary Figs. 7f-g**), ACE2 is restricted to the apical face of primarily ciliated and goblet cells, with some expression on club and basal cells^27–32^. DGE analysis (**Supplementary Data 1**) confirmed the statistically significant upregulation of *ACE2* and *TMPRRS2*, as well as that of other membrane proteases and toll-like receptors implicated in SARS-CoV-2 infection and epithelial cell responses (**Fig. 4e**)^32–34^.

**Fig. 4.**
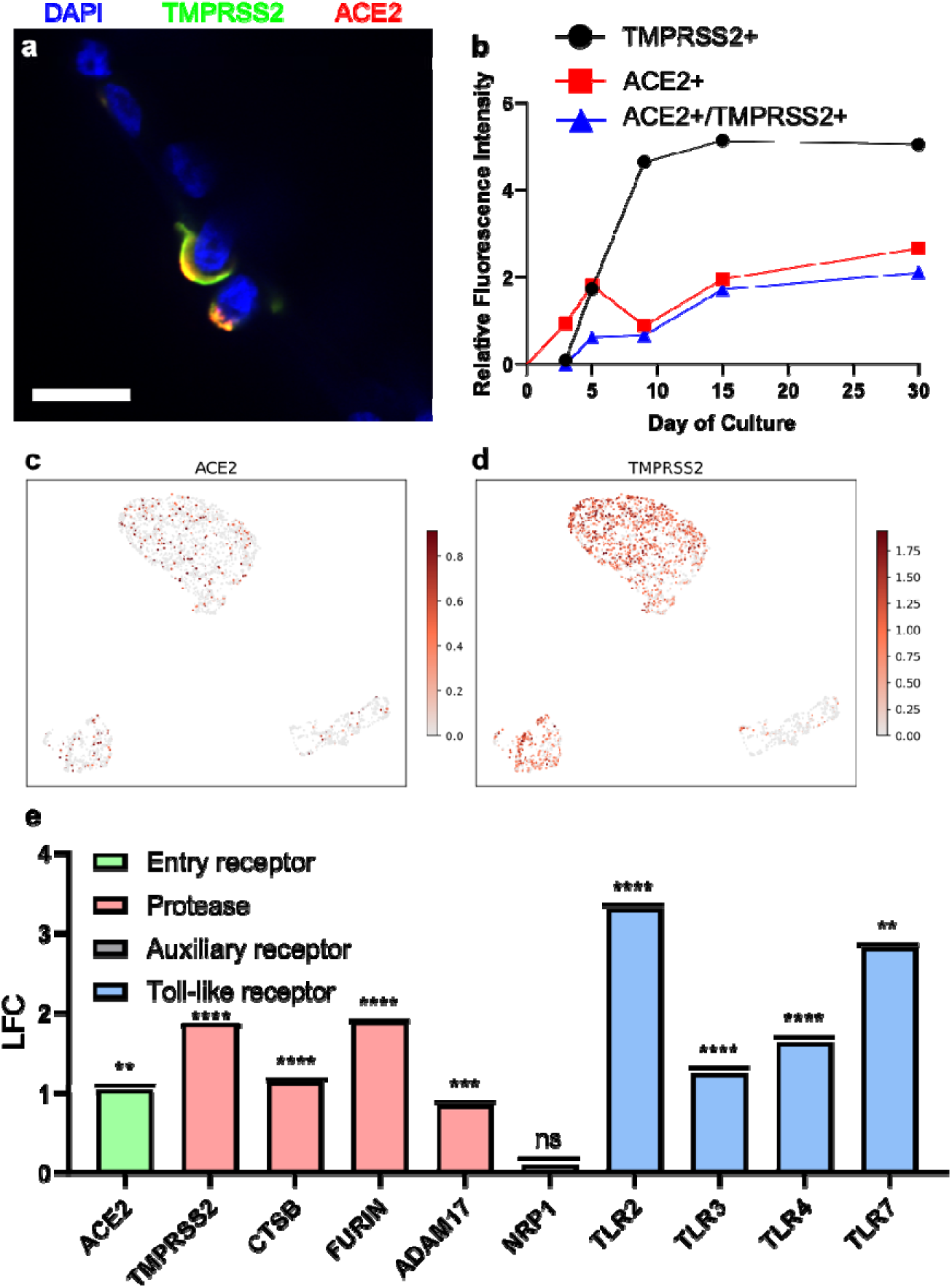
AORBs robustly express SARS-CoV-2-implicated entry factors. **a,** Immunofluorescence imaging confirms apical-out expression of ACE2 (red) and TMPRSS2 (green). **b**, Semi-quantitative analysis results show a time-dependent increase in levels of fluorescence for ACE2+, TMPRSS2+, and ACE2+/TMPRSS2+ cells, *n* = 6 AORBs per timepoint. **c**, scRNAseq *ACE2* expression. **d**, scRNAseq *TMPRSS2* expression. scRNAseq reveals higher *TMPRSS2* expression levels than *ACE2*. DGE analysis reveals upregulation of many other SARS-CoV-2 entry factors in AORBs with respect to 2D monolayer culture, *n* = 3 biological donors per experimental condition. Scale bar, 20 µm in **a**.

**Fig. 5.**
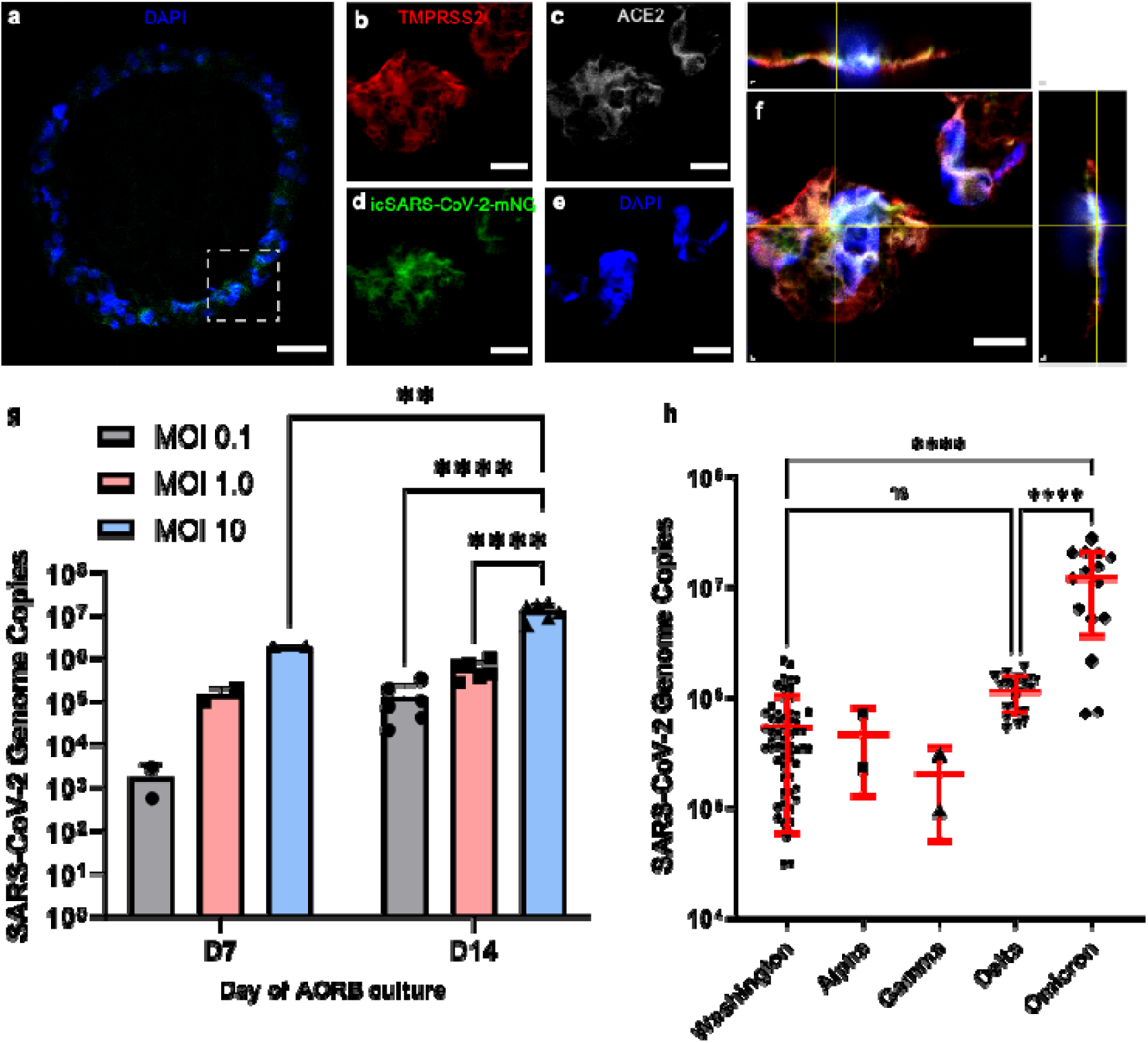
AORBs are infected by SARS-CoV-2. **a**, Section of a AORB overlay stained for nuclei (blue, DAPI) and replicated SARS-CoV-2 (green, icSARS-CoV-2-mNG). **b**-**f**, Zoomed in immunofluorescence images of stains for TMPRSS2, replicated viruses, ACE2, nuclei, and an overlay and orthogonal projections of these four markers, respectively. **g**, Increasing the MOI results in expected increases in replication. More mature AORBs show higher infectibility consistent with their higher expression of ACE2/TMRPRSS2 levels (see Fig 4b), *n* = 2 for D7 per MOI, *n* = 6 from 3 separate experiments in duplicates per MOI. **h**, AORB infection results with 5 different SARS-CoV-2 variants, *n* = 60, 2, 2, 21 and 15 for the five tested SARS-CoV-2 strains. Scale bars, 100 µm in **a**, and 10 µm in **b**-**f**.

### SARS-CoV-2 efficiently infects AORBs with >95% yield

Utilizing an infectious SARS-CoV-2-mNeon Green expressing clone (icSARS-CoV-2-mNG)^35^, we demonstrated replication of icSARS-CoV-2-mNG in AORBs (**Figs. 5a-f**), with the reporter detectable throughout the AORBs (**Fig. 5a)** and co-localized to cells co-expressing ACE2 and TMPRSS2 (**Fig. 5f**) at 72 hpi. Orthogonal projections taken at the depth of the nucleus showed intracellular expression of the mNG reporter (**Fig. 5f**), further validating SARS-CoV-2 infectibility. As expected, because ACE2 and TMPRSS2 expression is higher, the more mature Day-14 AORBs showed higher infectibility than Day-7 AORBs (**Fig. 5g**). RNAseq analysis confirmed expression of all SARS-CoV-2 genes in infected AORBs (**Supplementary Fig. 10a**). Viral yields were ∼0.5 log lower in AORBs compared with 2D and Transwell ALI cultures (**Supplementary Fig. 10b**). This finding is particularly impressive considering the lower initial virus amounts used to maintain a constant MOI. It is worthwhile noting that organoids formed from sub-optimal media conditions (**Supplementary Fig. 10c)** failed to support productive virus replication (**Supplementary Fig. 10b**), indicating the critical importance of the optimized SOP. An optimized infection of 72 h at MOI 1.0 resulted in high levels of infection (>95% of AORBs) and was used for subsequent assays.

We confirmed that not only the ancestral SARS-CoV-2 Lineage A Washington strain, but also Alpha, Gamma, Delta, and Omicron variants, all productively infected AORBs (**Fig. 5h**). At 3 dpi, SARS-CoV-2 genome copies were within the same order of magnitude for Washington, Alpha, and Gamma, while the genome copies for Delta and Omicron were 1–2 orders of magnitude higher with respect to the others (**Fig. 5h**), consistent with previous observations^36^. There was no significant difference in replication between donors for both Delta and Omicron (**Supplementary Fig. 10d**).

### RNAseq reveals differing epithelial responses to Washington, Delta, and Omicron

The top 10 upregulated biological processes found by GO analysis performed on AORBs infected with SARS-CoV-2 Washington strain vs. uninfected negative controls included ribosome processing, translation, and respiration (**Figs. 6a-b**). This was in direct contrast to the downregulated processes for AORBs with respect to 2D basal cells (**Fig. 2e**). SPIA identified three relevant activated pathways involving COVID-19, NF-κB and TNF signaling (**Fig. 6a**; **Supplementary Data 5**). Further analyses of subsets of relevant host epithelial defense response genes for Washington, Delta, and Omicron showed similar transcriptional changes upon infection with Washington and Omicron strains, and some difference in Delta compared to the other two (**Fig. 6c**). More specifically, Washington and Omicron strains showed similar patterns of transcriptional changes in type I interferon (IFN) (*ISG15*, *IRF3*, *IRF7*), cytokine (*TNF*, *IL1B, CXCL8*, *MMP1*, *MMP3*), NF-κB (*NFKB1, NFKBIA, RELA*), immune mediator (*CD86*, *CEACAM5*), and apoptosis (*CASP3*, *CASP7*, *CYCS*, *FADD*, *APAF1*) genes, with a few minor differences (*IRF7*, *IL6*) (**Fig. 6c**). The notable transcriptional differences observed with the Delta strain included quantitative changes in overlapping genes for Washington and Omicron with an increased extent of upregulation of other immune genes (*IFNL1*, *IFNG*, *IL12B*), and downregulation of certain apoptosis genes (*CASP3*, *CASP7*, *CYCS*) (**Fig. 6c**).

**Fig. 6.**
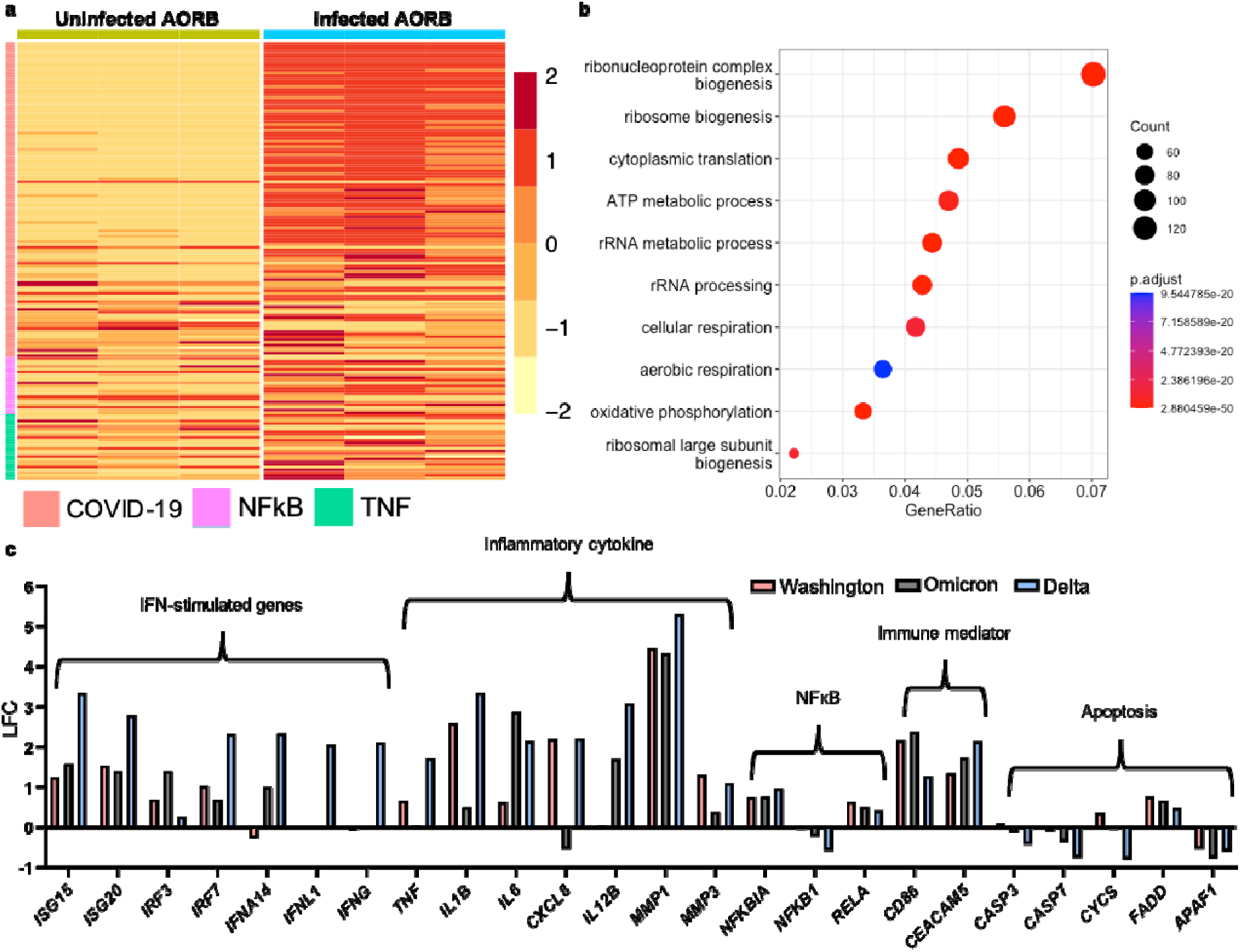
Transcriptomic analysis of infected AORBs. **a,** Heatmap summarizing three statistically significant activated pathways comparing infected AORB vs. uninfected control identified by SPIA (COVID-19, hsa05171; NFkB, hsa04064; TNF, hsa04668). **b**, Top 10 statistically significant upregulated biological processes from GO analysis of the enriched gene sets from DGE analysis. **c**, DGE analyses for 5 immune-mediator gene clusters comparing Washington-, Delta-, and Omicron-infected AORBs, respectively, showing different profiles of transcriptomic signatures in AORBs infected by the three strains, *n* = 3 biological donors per experimental condition.

### AORBs provide an efficient anti-SARS-CoV-2 compound HTS platform

For SARS-CoV-2, nucleoside analogs (NAs) remdesivir^37^ and molnupiravir^38^, and 3C-like protease inhibitor (PI) Paxlovid^39^, are the only FDA-approved antiviral regimens to date. We tested the antiviral potency of remdesivir, PF-07321332 (the PI component of Paxlovid; nirmatrelvir) as well as of a guanine double NA prodrug, AT-511, currently in Phase 3 trial (Bemnifosbuvir)^14,40^. Cell infection models (AORBs, hBTEC-ALI, Calu-3, Caco-2, and Vero) were treated with 0 - 10 μM of compound and infected with the Washington strain for 48 (Vero) or 72 h (AORBs, hBTEC-ALI, Calu-3, Caco-2). Mean effective concentrations (EC_50_) and EC_90_ values for all compounds were within range of published values (**Table 2**)^41–46^. AT-511 applied against Washington strain in these 2D cell lines has not been reported, but, in our hands exhibited limited activity. AT-511 is a double prodrug requiring multiple cellular enzymes for processing into its active nucleoside 5’-triphosphate form (NTP; AT-9010)^45^. We postulated that primary differentiated cultures might have an enhanced ability for AT-511 ➔ AT-9010 conversion over 2D cultures used in this study. To investigate this, cells were treated with 10 µM of AT-511 for 4 hr, then the cellular fraction collected for analysis. hBTEC cells had significantly higher NTP levels (170.8 ± 62.5 pmol/10^6^ cells) compared with Vero (0.89 ± 0.05 pmol/10^6^ cells), Calu-3 (4.93 ± 0.06 pmol/10^6^ cells), and Caco-2 (30.1 ± 2.4 pmol/10^6^ cells), which could be related to its greater potency in hBTEC cells (**Supplementary Table 2**).

Overall, EC_50/90_ were lower in the ALI model compared with AORBs; however, these differences were statistically insignificant. While AORB cultures had higher levels of variability compared with ALI cultures, much of this is attributed to donor variability (**Supplementary Table 3**). To investigate whether donor variability was an issue specific to AORBs, we generated matched donor ALI and AORB cultures (donor 3) and performed concurrent dose response assays which did not reveal any significant difference in obtained EC_50/90_ values (data not shown).

## Discussion

Historically, 3D cellular constructs with polarity reversal have been reported in human enteroids^47^, airway spheroids^48,49^, mammary acini^50,51^, and kidney cysts^52,53^. Generally, such polarity inversion is pathological^50,51,54^ and/or probabilistic^52,55,56^. Of these, it is important to recognize the early work by Pederson et al., where ion transport was demonstrated using apical-out nasal organoids, although the underlying mechanism of polarity inversion was not further studied^48,49^. Recent pioneering work from Amieva et al. developed the ECM removal methodology to reproducibly induce geometrical eversion of organoid polarity in a human enteroid model, producing apical-out morphology^57^. Additional studies have since adapted the ECM removal methodology, including cell-mediated ECM degradation, to other types of organoids^11,58–66^. Also, manipulation of the Wnt/β-catenin and BMT signaling pathway has been proposed^67^. Such geometrically-inverted polarity has facilitated studies of host-pathogen interactions^11,12,58,61,62,64,68^, ciliary dynamics^66^, barrier integrity^57,63^, nutrient uptake^57,60^, bio-inspired robots^69^, and more^59,65,66^. However, the ECM removal method of apical-out organoid cultures based on polarity eversion often results in undesirable instability and varying degrees of “apical-out-ness”, both spatially and temporally. Such impractical results could be attributed to the poor and/or partial degree of the everted morphology. Additionally, apical-out airway organoids tend to exhibit positively skewed ciliated cell populations, ranging from approximately 30% to 90%^12,58,66,68^. Such inadequate proportions of *in vivo*-like cell heterogeneity may be attributed to the lack of ECM signaling. Furthermore, because ECM removal methods rely on the conventional Matrigel embedding protocol, associated limitations, such as poor scalability, high variability, imaging/analysis challenges, and phenotype heterogeneity, remain unresolved^70^.

Extensive characterization confirmed that differentiated AORBs had stable apical-out polarity, physiologically relevant cell compositions and transcriptional profiles, and orientation with apical-faced beating cilia and mucus production. Apical-out orientation of secretory cells was confirmed with a functional assay involving IL13-induced goblet cell hyperplasia, where gel-like substances visible outside AORBs in bright-field images were confirmed to contain mucins with fluorescent lectin. Additionally, the upregulation of *CLCA1* and *ALOX15B* were consistent with previous *in vitro* studies on IL13-induced *MUC5AC* expression^26,71,72^, suggesting similar cellular effects despite the geometrically inverted morphology. Moreover, AORBs having a large diameter facilitated greater cell coverage per individual organoid. Sufficient RNA was extracted from a single organoid, and a passing QC score was achieved upon subsequent sequencing library preparation. GO analysis demonstrated that the top 5 upregulated biological processes involved signaling pathways for ECM organization, cell-cell adhesion, and epithelial tube morphogenesis, while the top 5 downregulated biological processes related to cellular transcription/translational machinery. An additional reduction in proliferative gene expressions supported evidence of a differentiated state of cells in AORBs. Lastly, scRNAseq confirmed physiologically representative cell types and their ratios, further validating AORBs. It is worth noting that, while the cellular heterogeneity of previous apical-out organoids has been somewhat studied^12,58,66,68^, AORBs were one of the first to date to be rigorously studied using scRNAseq^73^.

Since the onset of the COVID-19 pandemic, a plethora of airway organoid models for SARS-CoV-2 studies have been reported. The field has gained attention for its role in facilitating swift and transparent research efforts aimed at saving lives, despite some associated side effects. The results are highly variable in nature, frequently presenting contradictory outcomes, even within the same organ system^13^. Furthermore, there is a significant lack of information regarding infection yields. This deficiency can be attributed to two main factors: first, a rush for timely publication, where the importance of yields was often overlooked, and attention was solely directed toward viral replication in the models. Second, quantifying infection yield proves challenging in ECM-embedded organoids due to the limitations discussed earlier. Consequently, this makes it challenging to assess the practicality of these models^74–79^. Notably, in everted distal lung organoids^11^, SARS-CoV-2 yield is limited to <10%, and shows no viral tropism to ciliated cells, a primary target for SARS-CoV-2^80,81^.

The utility of AORBs was assayed as a platform for the study of SARS-CoV-2 and antiviral drug development. Robust expression of ACE2 (on the apical face) and TMPRSS2 in AORBs, as well as RNA-level expression of SARS-CoV-2 implicated entry factors, were confirmed. Unlike eversion organoids, AORBs were infected by SARS-CoV-2 in a high yield (>95% of all singular AORBs used). Efficient infection was attributed to the stable spatial and temporal apical-out characteristics of AORBs. The AORB model, with the one-per-well modularity with HTS capability, would provide a significant advantage in quantitation of single-organoid level quantitation of infection and antiviral response yields, hence promoting standardization through robust and rigorous evaluation.

Immune evasion by SARS-CoV-2 is an area of active study. Overall, our data support the prevailing literature that the antiviral response to SARS-CoV-2 infection in airway tissues is attenuated^82^. Of the three strains probed in detail, Delta induced a more robust antiviral response and production of inflammatory cytokines, including increased ISGs and NF-kB pathway components. The profile of Omicron-induced genes more closely mirrored that seen with the Washington strain, with the notable exception of inflammatory cytokines IL1B, IL6, and CXCL8. These data are consistent with recent observations^83,84^, though the difference in host antiviral responses to Washington versus other variants is still relatively understudied. The exhaustive RNAseq dataset provided herein may expand upon and facilitate such research endeavors. Another aspect worthwhile pointing out is the poor susceptibility of cell lines to Omicron variants^85^, failing to accurately represent human physiology. Our results corroborated with *ex vivo* data^36^, where Omicron variant in fact shows higher infectivity. Collectively, our results suggest that AORBs could provide an efficient and physiologically relevant platform to study different viral mechanisms such as immune evasion, virulence, and transmissibility.

The EC_50/90_ values determined in this study for remdesivir, AT-511, and nirmatrelvir were similar to published values. In the literature, Remdesivir potency in Vero cells ranges from 0.7 µM to over 20 µM^41–44,46,86^, while values in Calu-3 (0.01 µM – 2.4 µM) and Caco-2 (0.001 – 0.08 µM) cell lines are typically lower^41,42^. This discrepancy could be explained by differing level of enzymes required to process remdesivir to its active NTP form (e.g., carboxylesterase 1, CES1; and cathepsin A, CatA) in different cell types. Such enzymes are expressed at high levels in the lung^87,88^ and intestine^88,89^, which could underlie the improved activity of remdesivir in these cells. In ALI cultures, our EC_50_ values were comparable to published observations^42,46^, and not significantly different from those obtained in AORBs. Regarding antiviral efficacy of AT-511 in hBTECs, Good et al. reported an EC_90_ of 0.47 µM^45^, similar to Atea’s SUNRISE-3 clinical trial (∼0.78 µM)^14,40^. Our values from hBTEC-ALI and AORB cultures aligned with these reports. AT-511 had limited activity in 2D cultures. Similar to our 2D results, Do et al. reported no activity of AT-511 against a Beta variant strain in the Vero E6 clone and human hepatoma cells (Huh-7). However, in contrast to Good et al., the ATEA clinical trial, and our data herein, Do et al. did not observe activity in hBTEC-ALI cultures. They attributed this discrepancy to possible differences in metabolism of AT-511 to AT-9010 in the different cell culture systems and/or differences in the experimental approach. Our metabolism assay supports that 2D cell lines employed herein have a reduced capacity for metabolism of AT-511 compared with hBTEC-ALI cultures. Values obtained for nirmatrelvir were consistent with published observations of sub-micromolar activity in 2D and primary lung cell lines^90^.

With the FDA lifting the mandate for animal testing through the Modernization Act 2.0^91^ and further proposals advancing in the FDA Modernization Act 3.0^92^, microphysiological systems are gaining recognition for their potential as predictive preclinical models. In antiviral screening results, AORBs successfully detected true positive antiviral efficacy where 2D cell lines gave false negative results. It is worth noting that in drug development scenarios, false positive results in drug efficacy add unnecessary cost but will eventually be eliminated in later screening stages by lower throughput but more physiological tests (e.g., animal model, and clinical trials). False negative results, on the other hand, lead to irretrievable loss of lead compounds. Avoiding such loss requires assays that not only faithfully recapitulate human physiology, but also are sufficiently high-throughput to allow their use in the earlier stage screens. AORBs can help fill this unmet need for assays that are both physiologically relevant and high throughput. AORBs can also reflect donor variability, as evidenced by the varying replication potency and antiviral efficacy observed in AORBs derived from different donors. Because primary cells from donors are limited in quantity, the efficient use of cells (3000 cells per AORB; two orders of magnitude fewer than 24 well format Transwell cultures) afforded by AORBs is also enabling for future studies of biological variability. For example, just one P0 vial of cells can theoretically yield ∼ 210,000 AORBs. Furthermore, our minimal Matrigel scaffolding technique utilizes ∼2 orders of magnitude less ECM compared to the ECM-embedding method, reducing cost and increasing sustainability. Collectively, these findings suggest that AORBs will be a valuable *in-vivo-*like addition to the repertoire of preclinical models for drug development and disease modeling.

## Methods

### Chemistry

All compounds (remdesivir^93^, AT-511^94^, and nirmatrelvir^95^) were synthesized in-house by established protocols. Purity greater than 98% was confirmed by NMR and HPLC analysis.

### 2D immortalized cell line culture

For antiviral studies, the following immortalized/transformed cell lines were used: Caco-2 human colon epithelial cells (ATCC, HTB-37), Calu-3 human lung cancer cell line (ATCC, HTB-55), and Vero African Green Monkey kidney cells (ATCC, CCL-81). Media compositions were (1) Caco-2 and Calu-3: advanced Dulbecco’s modified Eagle medium:F12 (DMEM:F12), 10% FBS, 100 U/mL penicillin-streptomycin (pen-strep) (Gibco, 15140122), and 2 µM L-glutamine (L-glut), (2) Vero: DMEM, 10% FBS, 100 U/mL pen-strep. For all experiments, cells were grown at 37°C in a 95% O_2_, 5% CO_2_ incubator.

### 2D primary human bronchial epithelial basal cell expansion

Primary normal human tracheal/bronchial epithelial cells, hTBECs, (Lonza, CC-2540S) were expanded in Pneumacult^TM^-Ex Plus basal medium (STEMCELL Technologies, #05041) supplemented with Pneumacult^TM^-Ex Plus 50X supplement (STEMCELL Technologies, #05042) and 0.5 mL of 200 mM hydrocortisone using the manufacturer’s recommended seeding density of 3,500 cells/cm^2^ on a T-75 cell culture flask (Corning). The cells were subcultured at the confluence of 70 - 80%. Upon collection, cells were passaged, used for organoid culture, or cryopreserved. Cells beyond passage number 4 were not used, as they did not yield adequate organoid formation.

### Primary human bronchial Transwell air-liquid interface culture

For hBTEC-ALI cultures, 1.5×10^5^ cells were seeded on a 24-well 0.4 µm polystyrene transwell insert (Corning® cat. #3470) coated with type 1 bovine collagen type IV (Advance BioMatrix, PureCol) in complete PneumaCult™-Ex Plus medium supplemented with hydrocortisone. After 3 days, media was removed from the upper apical chamber to expose this surface to the air and complete Pneuma-Cult™-ALI medium supplemented with Pneuma-Cult™-ALI supplements (STEMCEL Technologies, #05001), hydrocortisone, and heparin solution added to the basal chamber to induce differentiation. Media was refreshed in the basal compartment every 2-3 days for 4 weeks. Culture viability and establishment of an intact polarized epithelial layer was routinely analyzed by microscopic analysis of cell morphology, cilia beat frequency, mucus formation and the absence of barrier-leakage. For all experiments, cells were grown at 37 °C in a 95% O_2_, 5% CO_2_ incubator.

### Preparation of hanging-drop plates

For at least one day prior to organoid seeding, a 384-well hanging-drop (HD) plate custom designed with injection molding (Xcentric) was soaked in 0.1% solution of Pluronic F108 (Sigma-Aldrich, 542342) in distilled water (diH_2_O) (Gibco, 15230). On the day of organoid seeding, the HD plate was rinsed off with diH_2_O and then air dried. The plate was then UV-sterilized (Analytik Jena, CL-1000) for 20 minutes on each side. While waiting, the wells of a sterile non-tissue-treated round-bottom 96-well plate (Corning, 351177) were each filled with 150 μl of diH_2_O supplemented with 1% pen-strep. When the HD plate was fully sterilized, it was placed (sandwiched) between the bottom and the lid of the 96-well plate. The two troughs of the short-edged sides were loaded with diH2O, which were then stuffed with sterile gauze pads. Lastly, the long edges were each added with ∼ 1 ml of diH_2_O.

### Hanging-drop AORB culture

The seeding solution was composed of Pneumacult^TM^ Airway Organoid Basal Medium (Stemcell Technologies, #05061) supplemented with the following additives: Pneumacult^TM^ Airway Organoid Seeding Supplement (Stemcell Technologies, #05062), methylcellulose (Sigma-Aldrich, 94378), and growth factor reduced Matrigel (Corning, 356231) (**Supplementary Table 1)**. First, the seeding supplement was added to the base medium (10% concentration). Next, an appropriate volume of methylcellulose was added (0.24% final concentration at the target volume) and the solution warmed in a 37°C bead bath (Fisher Scientific, GSGPD10). Once warm, Matrigel (at 5.35 μg/ml to 16.05 μg/ml final concentration) was then deliberately kept cold in ice and added to the pre-warmed media. Next, the desired volume of cell suspension at a final concentration of 120,000 cells/ml (3000 cells per 25 μl of the droplet) was added. Throughout the process, it was crucial to mix components very thoroughly. Immediately after the seeding solution was prepared, an eight-channel repeater pipette (Eppendorf, 4861000120) was used to load the HD plate in a staggered zigzag pattern (8 alternating rows x 24 columns = 192 wells). This pattern minimized adjacent wells with droplets from merging.

### Maintenance of AORBs

The organoids were monitored as needed with a high-throughput imaging system (Thermo Scientific, EVOS FL Auto 2). At days 3-4 of organoid seeding and every three days thereafter, the media exchange procedure was performed using a liquid handler robot (Analytik Jena, CyBio FeliX). Volumes to be removed from and added to the organoids were programmatically controlled using the CyBio Composer software. The organoid differentiation medium (ODM) to be exchanged with was Pneumacult^TM^ Airway Organoid Basal Medium (STEMCELL Technologies, #05061) supplemented with Differentiation Supplements (STEMCELL Technologies, #05063). It may be necessary to pre-fill wells around the outer edges with an additional medium (typically no more than ∼5 μl) to prevent aspiration of organoids. Moreover, it is advised to limit the time of handling HD plates outside the humidity-controlled incubator as much as possible to minimize evaporation of the liquid drops. The HD plate was kept in a cell culture incubator (Thermo Scientific, VIOS 160i) with 5% CO_2_ injection at 37°C.

### Virus preparation

The following reagents were deposited by the Centers for Disease Control and Prevention and obtained through BEI Resources, NIAID, NIH: SARS-Related Coronavirus 2, Isolate hCoV-19/USA-WA1/2020, NR-52281 (Washington strain) and Isolate USA/CA_CDC_5574/2020, NR-54011 (Alpha), The following reagent was obtained through BEI Resources, NIAID, NIH: SARS-Related Coronavirus 2, Isolate hCoV-19/USA/MD-HP01542/2021 (Lineage B.1.351), in Caco-3 cells, NR-55282, contributed by Andrew S. Pekosz. The following reagent was obtained through BEI Resources, NIAID, NIH: SARS-Related Coronavirus 2, Isolate hCoV-19/Japan/TY7-503/2021 (Brazil P.1), NR-54982, contributed by National Institute of Infectious Diseases. Delta and Omicron strains were a kind gift from Dr. Mehul Suthar. icSARS-CoV-2-mNG was a kind gift from Dr. Stefan Sarafianos. For Washington, icSARS-CoV-2-mNG, Alpha, Beta, and Gamma strains, virus stocks were prepared in Vero CL81 cells, while stocks for Delta and Omicron strains were prepared in Calu-3 cells. Virus titers were determined by ELIspot assay. In brief, Vero or Calu-3 cells were infected with 10-fold serially diluted stock in 2% infection media (base media with 2% FBS) for 90 minutes, after which inoculum was removed, and 2% methylcellulose supplemented with 2% FBS was added. After 2 (Vero) or 3 (Calu-3) days, cells were fixed with a 1:1 ratio of ice-cold methanol:acetone for 30 minutes for removal from the BSL-3 facility. Wells were blocked for 1 hour in Pierce™ clear milk blocking buffer (ThermoFisher, #37587), incubated with primary antibody against spike protein diluted 1:5,000 in blocking buffer (Elabscience, E-AB-V1006) overnight in a 37°C incubator with 5% CO_2_ injection, washed 3x with PBS, incubated for 1 hr with secondary antibody (HRP-conjugated goat anti-rabbit; Invitrogen, 656120) at room temperature, and washed 3x again then treated with 50 μl of True Blue™ peroxidase substrate (SeraCare, 5510-0030) for detection of viral antigens. Plates were read on a CTL plate reader (Immunospot®, ELISPOT reader). All virus infections were carried out in a BSL-3 level laboratory at Emory University in accordance with the guidelines of the 6th edition of Biosafety in Microbiological and Biomedical Laboratories and with the approval of the Emory University Environmental Health and Safety Office.

### Antiviral screening

For standard 2D antiviral screening, cells (Caco-2, Calu-3, and Vero) were grown to confluency (1×10^5^ cells) in 96-well plates. Dose-response curves were performed by treating cells with 2-fold serial dilutions (0 – 10 µM) of compounds of interest in respective base media containing 2% heat-inactivated FBS (ΔFBS) then infected with SARS-CoV-2 at an MOI of 0.1 (Vero) or 1.0 (MOI of 0.01 as reported previously^41^) (Caco2, Calu-3) for 48 (Vero) or 72 hrs (Caco-2, Calu-3). Cells/supernatants were collected in 150 µL RLT Buffer (Qiagen^©^) for downstream RNA extraction (Qiagen^©^, RNeasy 96 extraction kit) and subsequent RT-qPCR to detect viral load. For hBTEC-ALI, compounds were added at indicated dilutions to the basolateral chamber. Cells were washed 3x with HEPES-buffered salt solution (HBSS) on the apical surface to remove excess mucus, then infected by adding 50 µL of SARS-CoV-2 virus (MOI 1.0) to the apical chamber for a 5-hr adsorption after which virus was removed and cells retained in ALI for an additional 3 days. For AORB cultures, 3×10^3^ cells were seeded in hanging-drop suspension with Matrigel^®^ Basement Membrane Matrix (Corning, 356231) to generate a single organoid per well and cultured for 21+ days. AORBs cultured in the HD plate were first transferred to a 96-well ultra-low-attachment plate (Corning, 7007). Each well in the plate contains a single AORB. To achieve this, a P200 pipette tip with its entry edge cut using a sterilized razor blade is utilized to ensure transfer without shearing AORBs. Serially diluted compounds and virus (MOI 1.0) were added directly to the wells for a period of 3 days. AORB-infected cultures were collected in 150 µL RLT Buffer (Qiagen^©^), while hBTEC-ALI cultures were collected in 300 µL of Trizol™ Reagent and RNA extracted by phenyl-chloroform method according to manufacturers’ protocol (ThermoFisher Scientific). All experiments were performed three independent times, each including technical duplicates or triplicates.

### RNA extraction and RT-qPCR

RNA was detected by RT-qPCR using a 6-carboxyfluorescein (FAM)-labeled probe with primers against SARS-CoV-2 non-structural protein 3 (nsp3). (SARS-CoV-2 FWD: AGA AGA TTG GTT AGA TGA TGA TAG T; SARS-CoV-2 REV:TTC CAT CTC TAA TTG AGG TTG AAC C; SARS-CoV-2 Probe: 56-FAM/TC CTC ACT GCC GTC TTG TTG ACC A/3BHQ_1) RNA isolated from uninfected cells was used as a negative control for virus detection. RNA was added to optimized 10 µM primer/probe mix in Mastermix (Quantabio, qScript™ XLT One-Step RT-qPCR ToughMix^®^) and run on StepOne Plus real-time PCR (Roche) according to the manufacturer’s protocol. CT values were calculated from replicate groups, and virus yield was then quantified via standard curve.

### Cellular pharmacology assay

Cellular pharmacology assays were performed as previously described^41^. In brief, cells were plated and treated with 10 µM AT-511 for 4 hrs. Vero, Calu-3, and Caco-2 were seeded at 1×10^6^ cells per well in 12 well plates. hBTECs were grown at ALI with 1.5×10^5^ cells per transwell insert in 24-well plates. After 4-hr incubation with compound, cells were washed 2x with ice-cold PBS, then scrapped to collect the cell pellets in 70% methanol. Extracted samples were subjected to analysis by liquid chromatography mass spectrometry (LC-MS).

### AORBs cryosectioning

AORBs (15-20 per tube) were first pooled from the HD plate with a cut P200 tip to prevent shearing damage to AORBs. The medium was removed to the minimum extent possible. AORBs were then washed three times with 1X PBS (Gibco, 10010), with 2-3 minutes between washes. Next, AORBs were fixed in 4% PFA (Thermo Scientific, J61899-AP) in PBS for 30 minutes, followed by PBS washing three times. At this point, the fixed AORBs can be stored at 4°C or processed further with embedding procedures. The first step was to incubate AORBs with 0.5% methylene blue (RICCA, 4850-16) in PBS for ∼10 minutes. Next, AORBs were carefully and repeatedly washed until no blue stain bled out. AORBs were then transferred to optimal cutting temperature (Tissue-Tek, OCT) compound (Sakura, 4583) in a cryomold (Polysciences, 18986-1), with as little PBS as possible. The samples were flash-frozen by manipulating the cryomolds in partially frozen molecular biology grade isopentane (Sigma-Aldrich, M32631). The frozen blocks of samples may be kept at -80°C for long-term storage. A cryostat (Thermo Scientific, CryoStar NX70) at the histology core of the Parker H. Petit Institute for Bioengineering and Bioscience, Georgia Tech, was used for cryosectioning. Sections of 10-20 mm thickness were collected, and two to three sections were applied per glass slide. The slides were kept at -20°C until further use in hematoxylin and eosin (H&E) or immunofluorescence staining.

### Hematoxylin and Eosin staining and imaging

H&E staining was performed using an autostainer system (Leica, ST5010). Briefly, slides with AORB sections were first brought up to room temperature, and then stained by a pre-established protocol. The stained slides were then coverslipped with Cytoseal 60 (Richard-Allan Scientific) and imaged in color using a Leica DMi8 microscope.

### Immunofluorescence staining of cryosections

The slides were first brought to room temperature, and the boundary containing AORB sections was highlighted with a hydrophobic pen (Electron Microscopy Sciences, #71310) to minimize reagent use. This was followed by PBS washing 3x, permeabilization with 0.2% Triton X-100 (Sigma-Aldrich, T8787) in PBS for 5 min, PBS washing 3x, blocking with 4% BSA (Millipore Sigma, #82-067) for 1 h at room temperature, and an overnight incubation with primary antibodies at 4°C. The following primary antibodies were prepared in 1% BSA in PBS at the specified dilutions shown:

mouse anti-acetylated-α-tubulin (Santa Cruz Biotechnology, sc-23950; 1:50)

mouse anti-MUC5AC (Santa Cruz Biotechnology, sc-21701; 1:50)

rabbit anti-KRT5 (Abcam, ab53121; 1:1000)

rabbit anti-E-cad (Cell Signaling Technology, 3195S; 1:1600)

rat anti-ITGα6 (Santa Cruz Biotechnology, sc-19622; 1:50)

mouse Anti-ITGβ4 (ThermoFisher, MA5-17104; 1:100)

goat anti-ACE2 (R&D Systems, AF933; 1:500)

mouse anti-TMPRSS2 (Santa Cruz Biotechnology, sc-515727; 1:50)

rabbit anti-LAM5 (Abcam, ab14509; 1:200)

After overnight incubation, slides were washed 3x with 1% BSA solution, then incubated with secondary antibody solution at room temperature for 2 h. Appropriate combinations of conjugated secondary antibodies were used to match the host of the primary antibodies, and these included:

Alexa Fluor 488 donkey anti-goat (Invitrogen, A11055)

Alexa Fluor 488 donkey anti-rat (Invitrogen, A21208)

Alexa Fluor 568 anti-mouse (Invitrogen, A10037)

Alexa Fluor 647 anti-rabbit (Invitrogen, A31573)

### Immunofluorescence staining of whole-mount AORBs

Whole-mount immunofluorescence staining of AORBs was adopted from a previously established protocol^96^. Briefly, fixed AORBs were permeabilized with 0.1% Tween (Sigma-Aldrich, P9416) in PBS for 10 mins in 4°C, washed 3x with PBS, incubated with organoid washing buffer (OWB) composed of filtered (Millipore Sigma, SCGP00525) 0.1% Triton X-100 (Sigma-Aldrich, T8787) and 0.2% BSA (Millipore Sigma, 82-067) in PBS, at 4°C for 15 mins, then incubated overnight with primary antibody solution in OWB on a gentle rocker in 4°C. After the overnight incubation period with primary antibodies, AORBs were washed 3x with OWB with an incubation period of at least 20 mins between the three washing steps. The washed AORBs were then incubated with secondary antibody solutions prepared in BWB and rocked overnight at 4°C. Tubes from this step onward were wrapped with aluminum foil to prevent photobleaching. Upon finishing the incubation with secondary antibodies, AORBs were washed 3x with PBS, then counterstained with DAPI for ∼30 min at room temperature on a rocker. Counterstained AORBs were finally washed with PBS 3 times, and transferred either into a chambered coverslip (ibidi, #80821) or onto a coverslip depending on the magnification required. Samples were imaged on a Nikon W1 spinning disk confocal microscope.

### AORB IL-13 treatment

During the second media exchange routine, fresh medium with 1 ng/mL of IL-13 (Peprotech, #200-13) was used to treat AORBs. Treatment conditions were controlled to test both negative controls and IL-13 groups within the same plate. AORBs were imaged on EVOS to monitor morphology and secretions daily.

### Fluorescent lectin assay

Lectin with FITC conjugate (MilliporeSigma, L4895) was used for the fluorescence mucin binding assay. Manufacturer recommended concentration of the reconstituted lectin was added. AORBs were pooled into a 12-well plate, then imaged under a DMi8 microscope. NucBlue (ThermoFisher, R37605) was also used to track live cells along with lectin-bound mucin components.

### Bulk RNA sequencing

For bulk RNA sequencing, about five organoids per condition were harvested from the HD plate to DNase- and RNase-free tubes (Eppendorf, 022363344). In the case of 2D cells, adherent cells were collected with a cell scraper (Sarstedt, 83.1380). The cells/organoids were lysed in RLT lysis buffer (Qiagen, 79216) containing 1.0% β-mercaptoethanol (Sigma-Aldrich, M3148). The lysates were transported in dry ice to Emory Integrated Genomics Core (EIGC), Emory University, where further sample processing and sequencing were carried out. RNA was isolated using the Quick-RNA Microprep Kit (Zymo Research). The cDNA was generated using the SMART-Seq® v4 Ultra® Low Input RNA Kit (Takara Bio). The final sequencing library was made using the Nextera XT kit (Illumina). The library construction used Nextera XT chemistry, which tagmented 200 pg of DNA and added adapters in one step. The fragmented and tagged DNA was then amplified for 12 cycles to incorporate dual-indexing sequencing adapters to generate the final library. A combination of both concentration and library size patterning was used as QC metrics to determine the success of the assay. UV Qubit fluorometer quantitation was used to determine the concentration of the library. A Bio/Fragment Analyzer trace revealed the size distribution and concentration of the final sequencing library. Based on these three parameters, we assigned library scores to each library. Those with a passing score were then used for sequencing in NovaSeq (Illumina). The read depth for the run was ∼40 million reads/sample.

### Differential gene expression and functional analysis

Demultiplexed raw FASTQ files provided by EIGC were downloaded to Georgia Tech’s high-performance computing cluster “Partnership for an Advanced Computing Environment (PACE)”. Quality control was first performed using FastQC^97^; *trimmomatic* was then used to trim the raw FASTQ files^98^. Upon passing quality requirements, the read fragments were aligned to the reference human genome (GRCh38.p13) or SARS-CoV-2 genome (GENCODE) using the STAR algorithm^99^. A custom Python script was written to generate a gene count matrix. Principal component analysis was performed to sort out any outliers. Gene-level differential gene expression (DGE) analysis was conducted with DESeq2 package in R^100^. Briefly, raw reads were normalised by the DESeq2’s median of ratios method. Identified outliers and genes with low counts (mean counts < 2) were filtered out and not included for subsequent hypothesis testing.

Lastly, computed log2fold change (LFC) values were shrunken with the ashr package to obtain more accurate estimates of LFC^101^. Differential expression testing was performed using the Wald test with Benjamin-Hochberg multiple comparison correction. The cutoff for significance was set at an adjusted p-value <0.05 and LFC of ±0.6. Heatmaps and volcano plots were generated using the ggplot2 package. Gene ontology (GO) and gene set enrichment analysis (GSEA) was conducted in clusterprofiler^102^, and pathway analysis was performed by signalling pathway impact analysis (SPIA)^103^.

### scRNAseq

scRNAseq of AORBs used the PIPseqTM pipeline (PIPseqTM T2 3’ Single-Cell RNA kit v4.0, Fluent BioSciences). The protocol was adapted from previous work^104^. Briefly, AORBs were harvested onto a cell strainer (431750, Corning) to remove debris. The AORBs were washed and pooled into a 1.5-mL Eppendorf tube, then dissociated into a single-cell suspension by incubating in TrypLE (12604021, Gibco) at 37°C for 30 min with frequent mixing by pipetting. Subsequently, cell viability was enhanced using a dead cell removal kit (130-090-101, Miltenyi Biotec). The resulting purified cell suspension with improved viability (>95%) was diluted according to the manufacturer’s recommendation (1250 cells/µL). PIPSeq kits were applied to capture mRNA, synthesize cDNA, and construct libraries. Libraries that passed QC were then used for sequencing on a 100-cycle NovaSeq SP kit, with read lengths as specified by the user guide. FASTQ files were first preprocessed with the same QC and trimming tools as described above. Subsequent computational analysis of the raw data was performed using PIPseeker v3.0 Software. RNA data processing steps included mapping with STAR aligner, transcript counting, cell calling, clustering, and differential expression. UMAPs and feature plots were generated as the final outputs.

### Statistical analyses

GraphPad Prism 10.0.2 was used to perform statistical analyses. Median effective concentration of compounds (EC_50_) and concentrations with a 90% inhibitory effect (EC_90_) of the three compounds tested were calculated by four parameter logistic curve regression and reported as the mean ± standard deviation. One-way ANOVA and subsequent Tukey’s post-hoc tests were used to compare the means of AORB diameters at differing Matrigel levels, donors, and batches. An alpha of 0.05 was used as a threshold for statistical significance.

## Data Availability

The main data supporting the results in this study are available within the paper and its Supplementary Information. Custom python and R scripts are available upon request. The raw and analyzed datasets are available for research purposes from the corresponding author upon reasonable request. Our RNAseq data have been deposited on Gene Expression Omnibus of National Center for Biotechnology Information and are available under the SuperSeries accession number GSE249581.

## Acknowledgements

This research was supported by NIH (R01-HL-136141; R01-AI-161570; R01-HL-139876), NSF EBICS (CBET 0939511), Price Gilbert Jr Chair’s Fund, Biomedical Engineering Department Seed funds, and the Center for AIDS Research at Emory University (P30-AI-050409). The study was also supported in part by the Emory Integrated Genomics Core (EIGC), which is subsidized by the Emory University School of Medicine and is one of the Emory Integrated Core Facilities. Furthermore, this work was supported in part through research cyberinfrastructure resources and services provided by the Partnership for an Advanced Computing Environment (PACE) at the Georgia Institute of Technology, Atlanta, Georgia, USA. We also wish to acknowledge the core facilities at the Parker H. Petit Institute for Bioengineering and Bioscience at the Georgia Institute of Technology for the use of their shared equipment, services, and expertise. We thank Dr. Greg Gibson and the Atlanta Single-cell Omics and Analytics Institute (AScOmAI) for assistance with scRNAseq. We also would like to thank Drs. Mehul Suthar and Stefan Sarafianos of Emory University for their kind donation of SARS-CoV-2 variants.

## Author Contributions

Conceptualization: Lee, Parigoris, Schinazi, Takayama; Formal analysis: Lee, LeCher, Shinagawa, Sentosa; Funding acquisition: Schinazi, Takayama; Investigation: Lee, LeCher; Methodology: Lee, LeCher, Manfredi, Goh, De, Tao, Schinazi, Takayama; Project administration: Lee, LeCher; Experimental approach: Lee, LeCher, Zandi, Amblard, Sorscher, Spence, Tirouvanziam, Schinazi, Takayama; Supervision: Schinazi, Takayama; Writing manuscript: Lee, LeCher, Takayama; Manuscript review & editing: all authors.

Lee and LeCher contributed equally to this work.

## Additional Information

**Supplementary information and data** are available for this paper at ∼

**Correspondence and requests for materials** should be addressed to Raymond F. Schinazi and Shuichi Takayama

**Peer review information** *Nature Communications* thanks the anonymous reviewers for their contribution to the peer review of this work. A peer review file is available.

---

**Supplementary Fig. 1.**
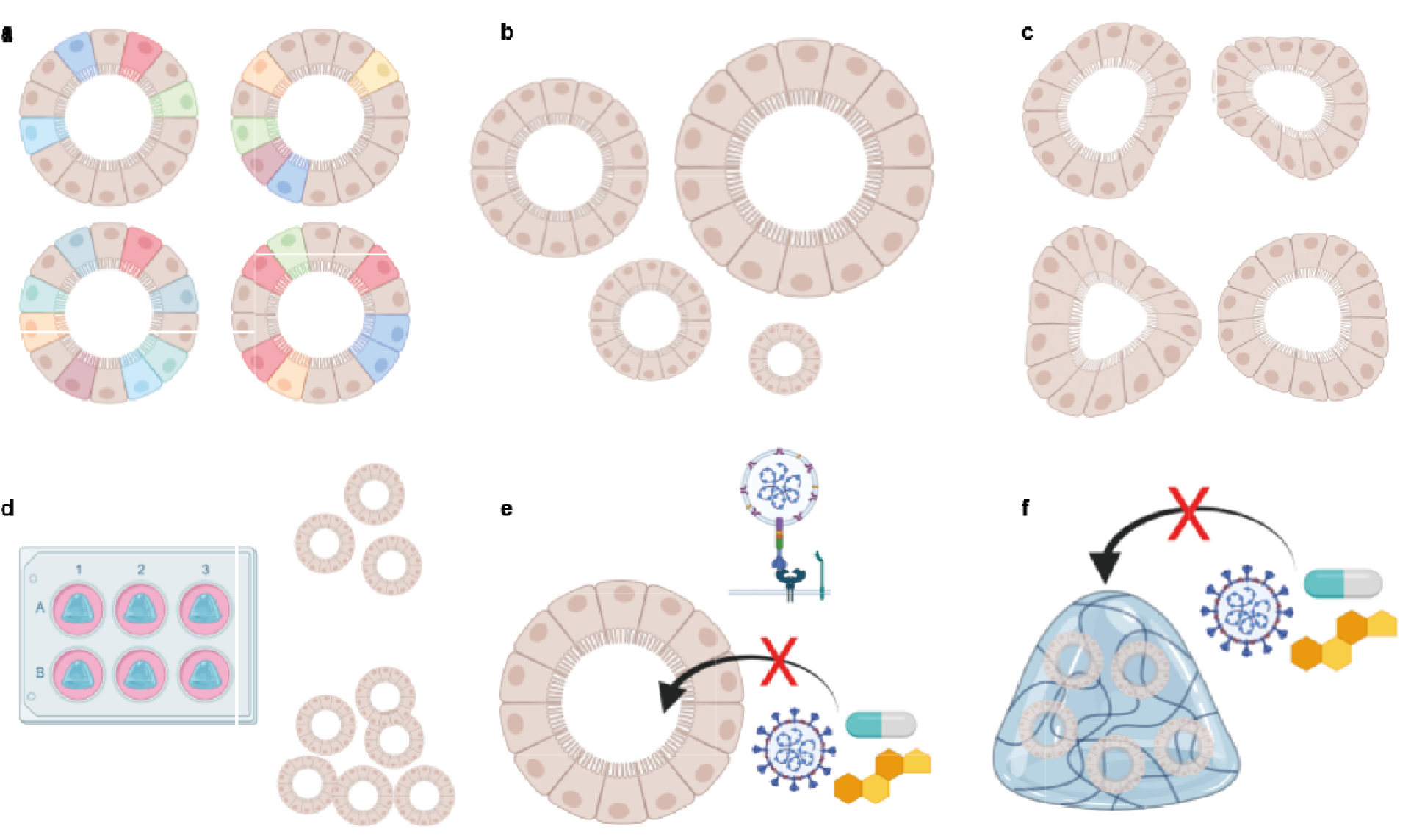
Limitations of ECM-embedded organoid culture. **a**, Difficulty controlling cellular heterogeneity of organoids embedded in ECM dome. **b**, High variability in organoid size. **c**, High variability in organoid shape. **d**, Variable numbers of organoids per well. **e**, Apical surface is sequestered away, hence preventing interactions with pathogen or compound of interest. **f**, ECM dome prevents efficient diffusion of pathogen or compound of interest.

**Supplementary Fig. 2.**
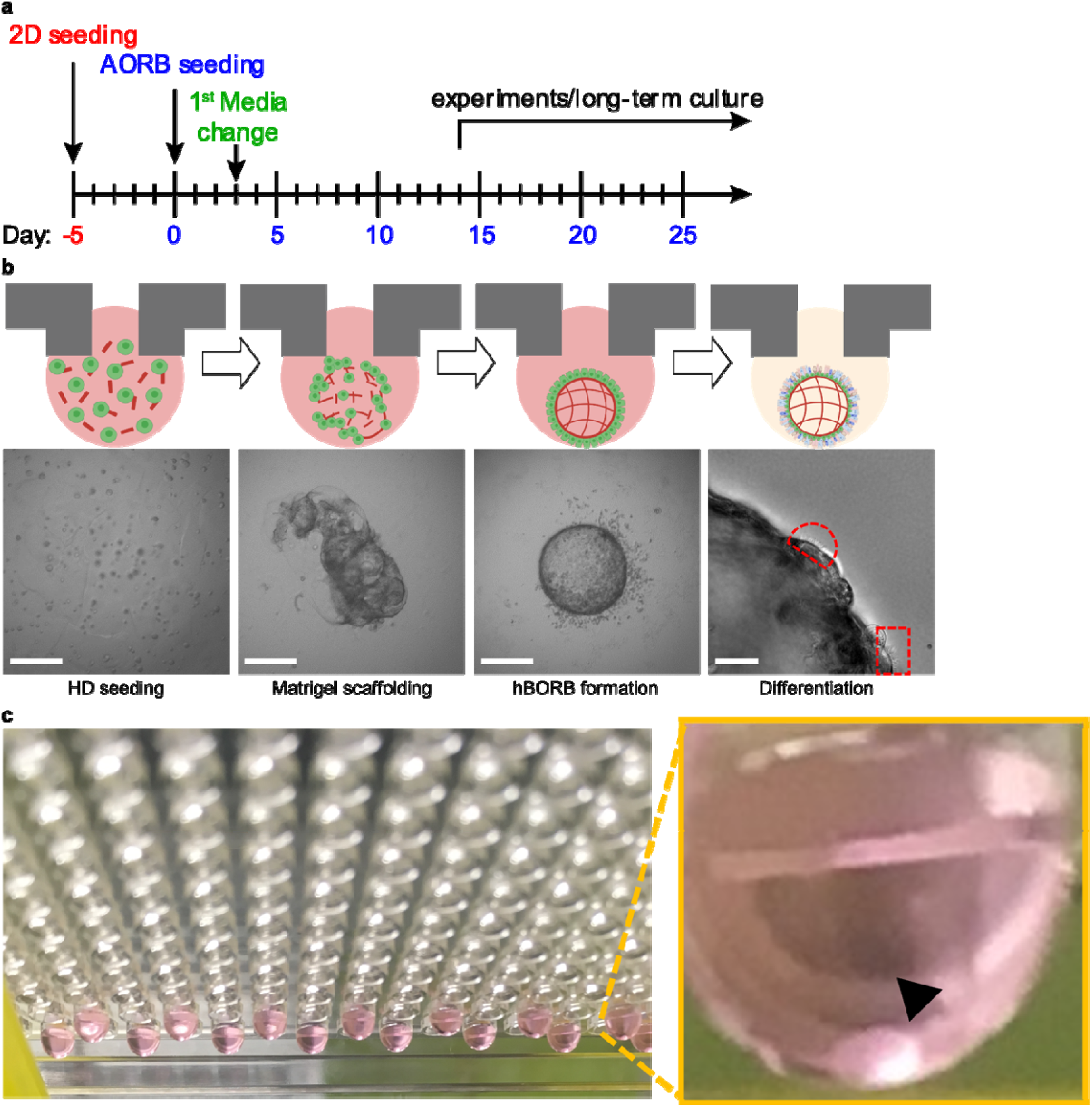
AORB culture overview. **a**, Time course summary of AORB culture. **b**, Schematic illustration and representative bright-field images in various phases of AORB culture (left to right, Day 0, 1, 4, 28). **c**, Photo of HD arrays and visual zoom of a singular AORB in one drop. Scale bars, 500 µm and 25 µm for the first three and right-most images, respectively.

**Supplementary Fig. 3.**
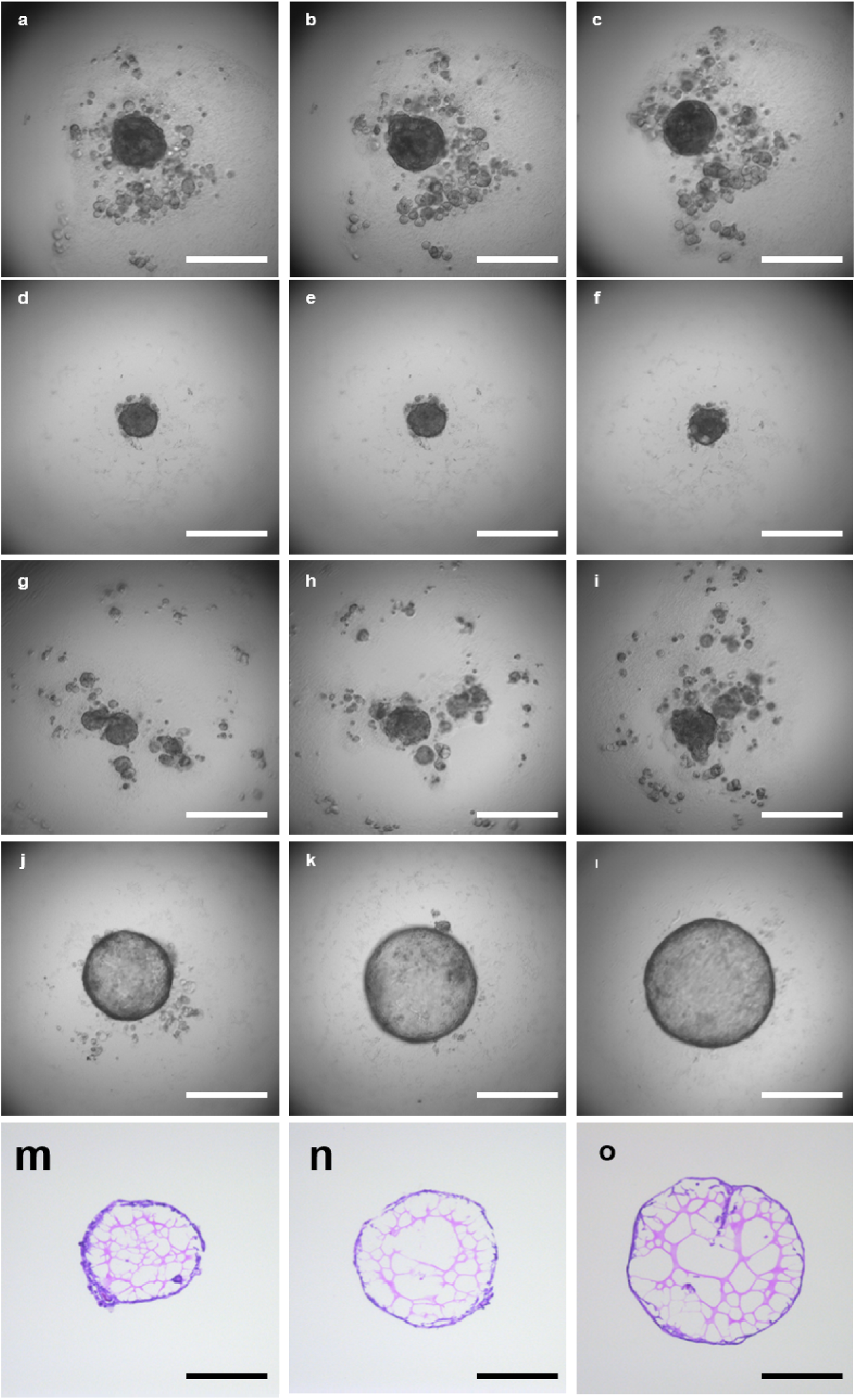
AORB culture optimization. **a**-**c**, 10% FBS+ condition yields multiple spheroid-like structures. **d**-**f**, Complete serum-free condition yields a singular small spheroid-like structure. **g**-**i**, Sub-optimal Matrigel also leads to improper formation of organoids. **j**-**l**, Increasing Matrigel amount within the working range (1300 – 2700 ng per 25 µl drop) augments size of AORBs. **m**-**o**, H&E staining confirms an acellular hollow core filled with Matrigel and a sheet of epithelial cells around the AORB boundary. Scale bars, 500 µm.

**Supplementary Fig. 4.**
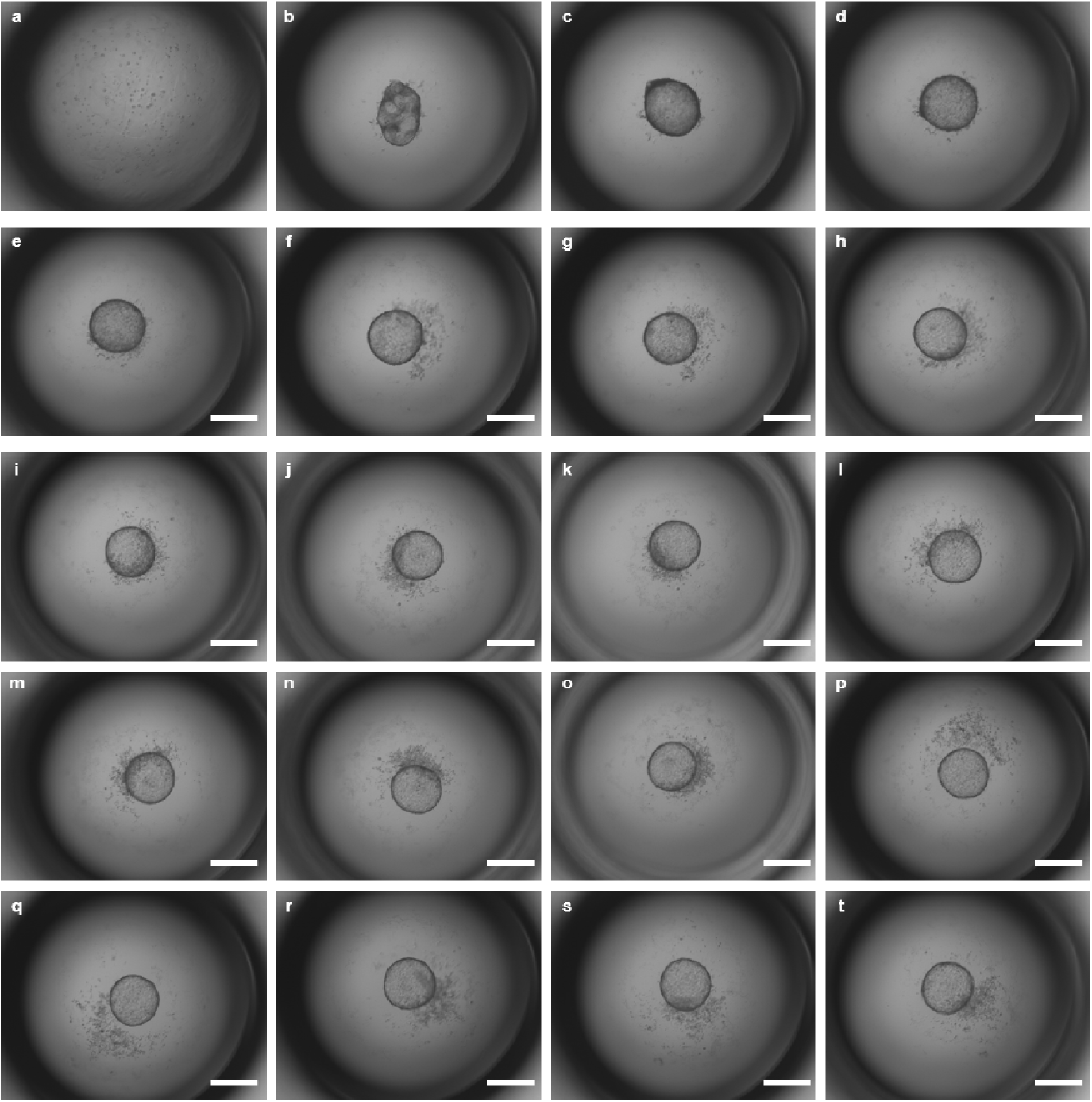
60-day AORB culture. **a**-**t**, Bright-field images of the same AORB at Days 0, 2, 4, 6, 25, 27, 29, 33, 36, 39, 40, 44, 48, 49, 51, 53, 55, 57, 58, and 60. Scale bars, 500 µm.

**Supplementary Fig. 5.**
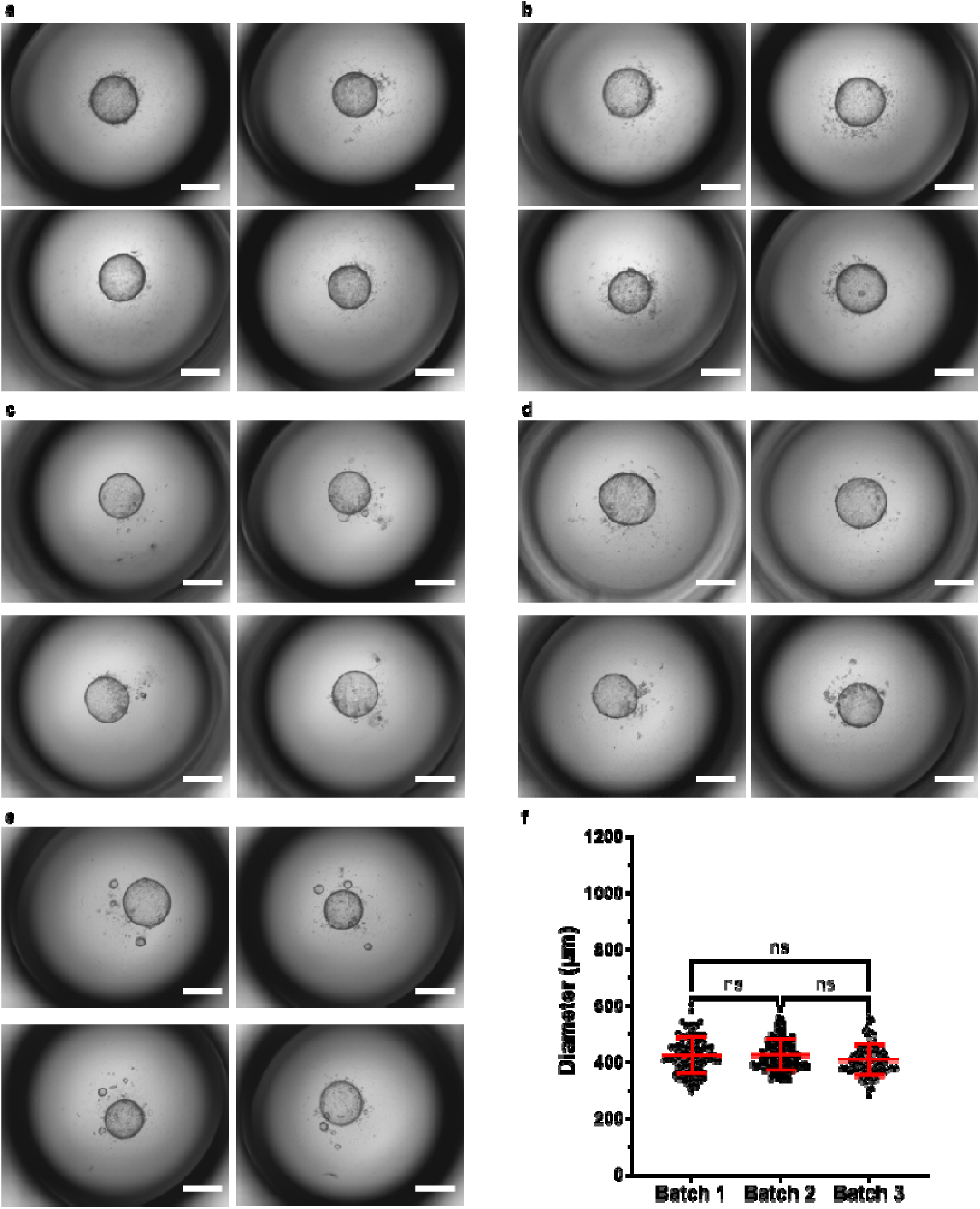
Biological and technical replicates. **a**-**e**, Bright-field images of AORBs derived from 5 different donors with 4 technical replicates each, respectively. **f**, There are no statistical differences in AORB diameter among distinct experimental batches (within donor 1), *n* = 96 for all three batches. Scale bars, 500 µm.

**Supplementary Fig. 6.**
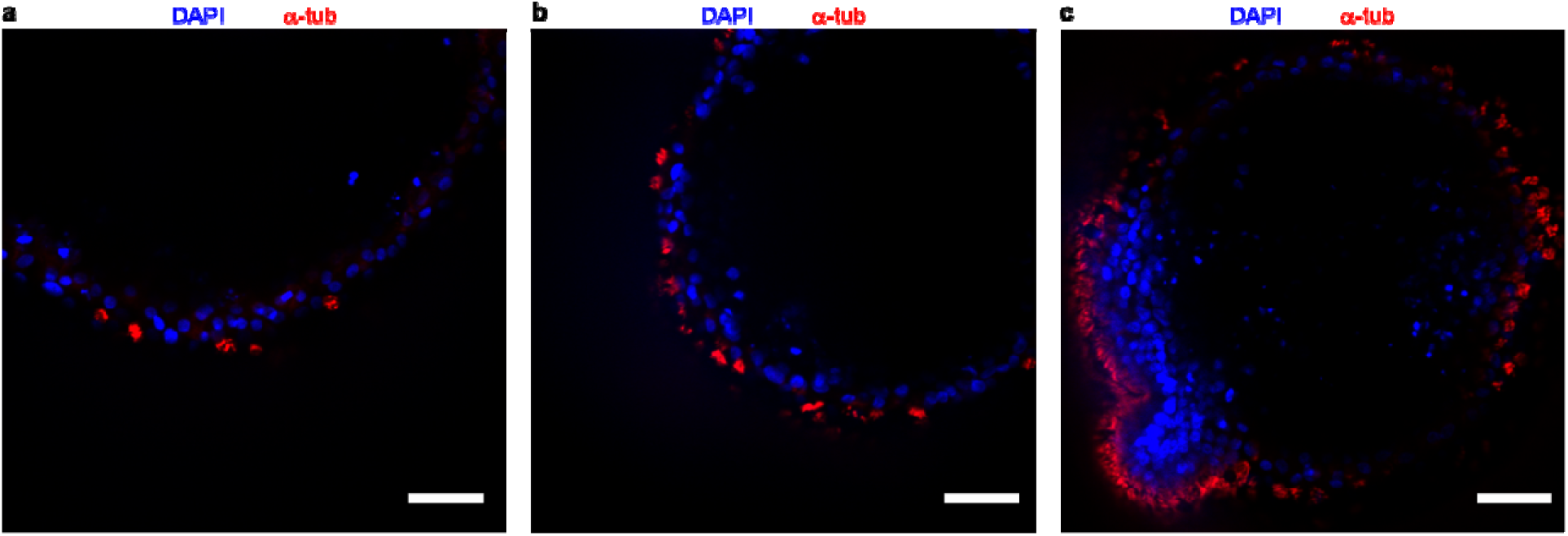
Time-course α-tub expression. **a**-**c**, Increasing fluorescence levels of α-tub AORBs are shown at days 14, 28, and 41, respectively. Scale bars, 100 µm in **a**-**c.**

**Supplementary Fig. 7.**
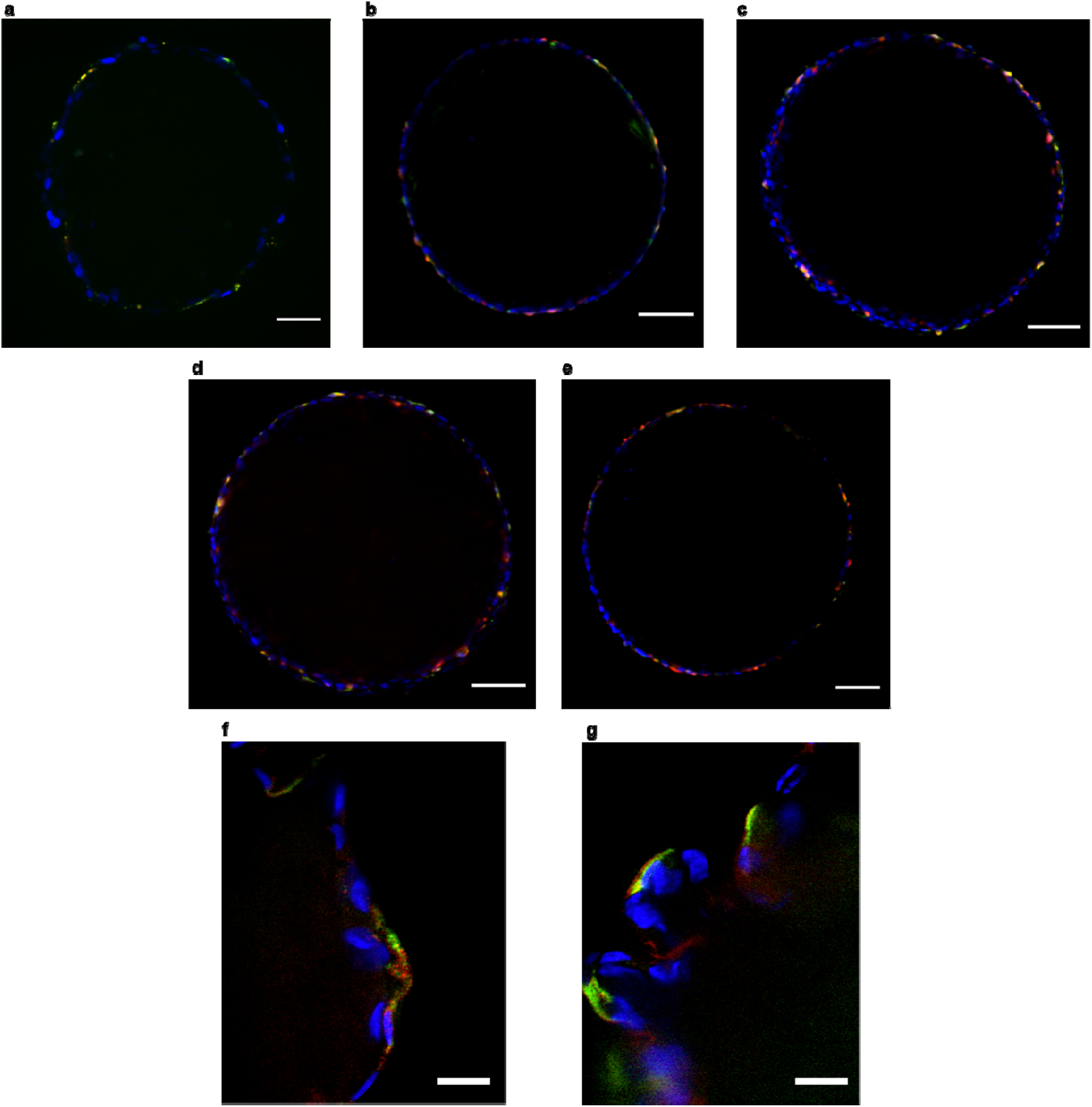
Additional ACE2/TMPRSS2 Expression Results. **a-e**, Confocal slice of AORBs stained for ACE2 (green) and TMPRSS2 (red) at days 4, 5, 9, 15, and 30, respectively. TMPRSS2 expression increases more dramatically than ACE2. High magnification images show apical localization of ACE2 (green) and TMPRSS2 (red). Scale bars, 100 µm in **a**-**e**, and 10 µm in **f**-**g**.

**Supplementary Fig. 8.**
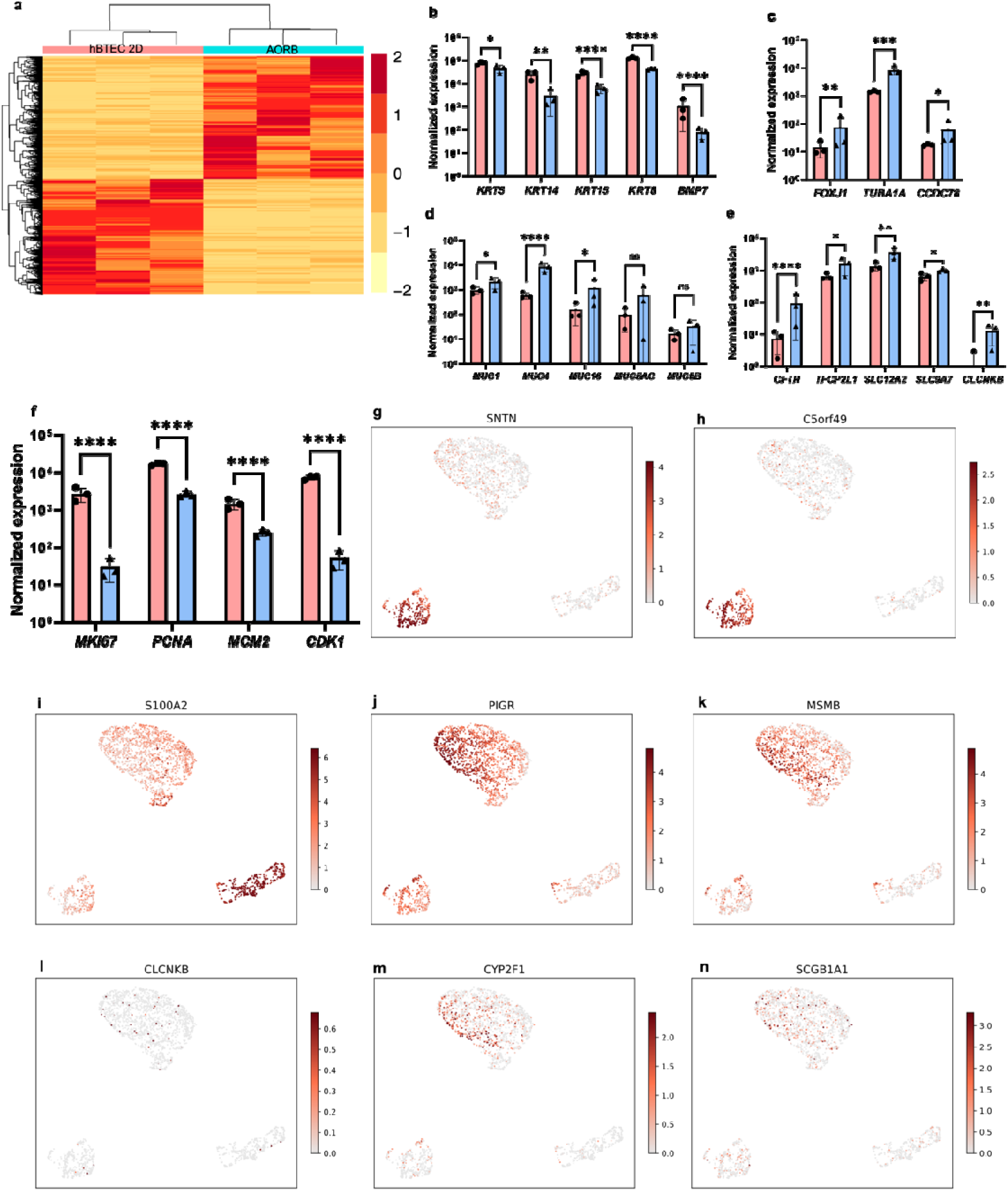
Extended results from bulkRNAseq analysis. **a**, Heatmap showing all statistically different genes clustered based on patterns of expression level (pink, 2D monolayer hBTEC culture; cyan, AORBs). **b**, Normalized expression counts for basal cell marker genes decrease in AORBs (blue) relative to their 2D culture counterparts (pink). **c**, Ciliated cell markers were upregulated in AORBs (blue) compared to 2D (pink). **d**, Normalized expression counts for membrane-associated mucins were higher in AORBs (blue) than 2D (pink), whereas canonical goblet cell markers were not statistically different between the two groups. **e**, AORBs exhibit upregulated pulmonary ionocyte markers. **f**, Bulk RNAseq normalized counts for 2D (pink) and AORB (blue) gene markers for proliferation. **g-m**, Expression of marker genes a feature plots on UMAP (**g-h**, ciliated; **i**, basal; **j-k**, goblet; **l-n**, ionocyte). *n* = 3 biological donors per experimental condition.

**Supplementary Fig. 9.**
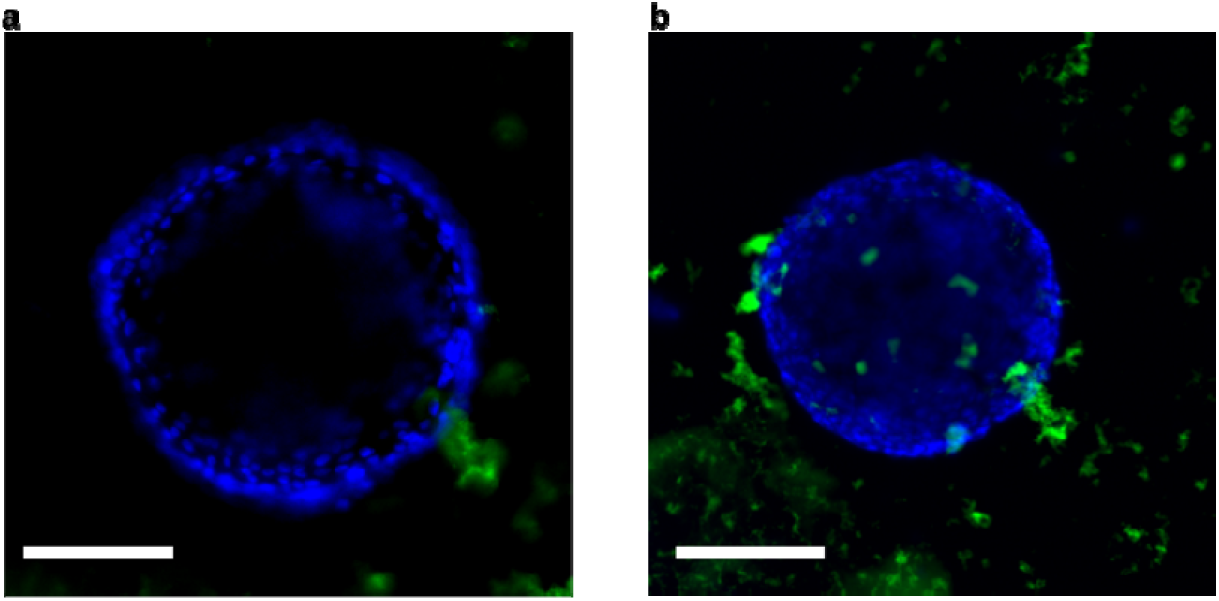
IL-13-treated AORBs. **a**-**b**, Immunofluorescence images of AORBs counterstained with DAPI (blue) and mucin stained with lectin fluorescent conjugates (green). Scale bars, 250 µm in **a** and **b**.

**Supplementary Fig. 10.**
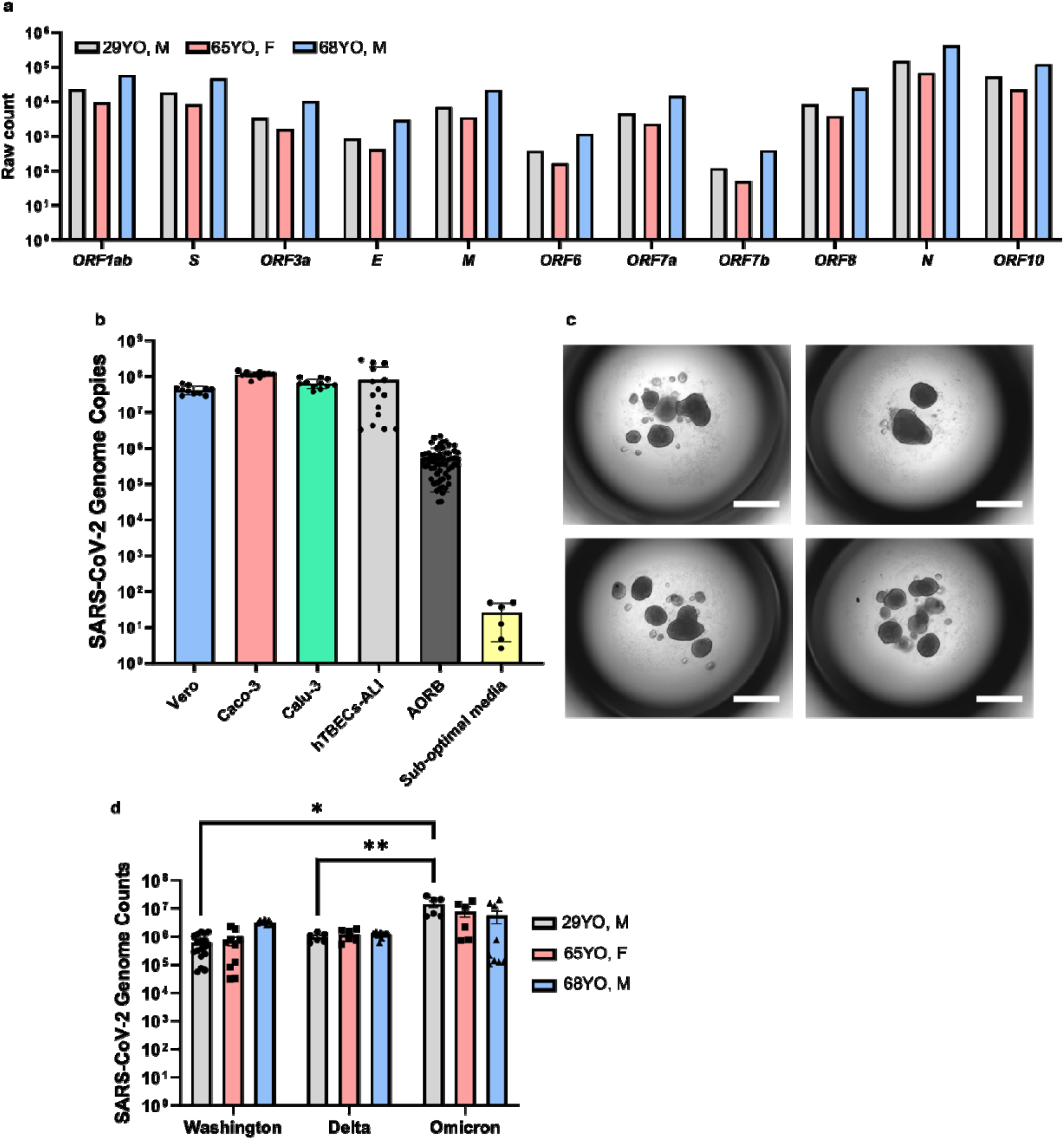
AORBs are a robust model for SARS-CoV-2 infection studies. **a**, SARS-CoV-2 genome mappings within Washington-infected AORBs exhibit all viral genes in RNAseq analysis, *n* = 3 biological donors per experimental condition. No viral genes are mapped from uninfected controls. **b**, AORBs show comparable infectibility to other established models, *n* = 12, 2D cell lines; *n* = 15 hTBECs-ALI; *n* = 60, AORB; *n* = 6 sub-optimal medium. **c**, Sub-optimal media conditions cause improper formation of organoids and do not support viral replication within them. **d**, Omicron variant yields higher infectibility than Delta and Washington across 3 donors, *n* = 18, 10, 8 for three donors, Washington; *n* = 6, 6, 9 for the three donors, Delta; *n* = 6, 6, 9 for the three donors, Omicron. Within variants, there is no statistically significant difference in replication among 3 donors. Scale bars, 500 µm in **c**.

