## Supplementary Information for "Development of robust antiviral assays using relevant apical-out human airway organoids"

Organoid, apical-out, SARS-CoV-2, antiviral testing, remdesivir, AT-511, nirmatrelvir, standardization, high-throughput

Experiments on Matrigel gelling in hBTEC media reconfirmed the necessity of adding cold Matrigel to prewarmed (37°C) seeding media (data not shown), consistent with previous findings^1–4^. Our reports included fetal bovine serum (FBS) as an empirically-identified gelling enhancer^1–4^. However, conditions with 10% FBS resulted in irregularly shaped spheroid-like structures (**Supplementary Figs. 4a-c**), while serum elimination led to the formation of a singular spheroid-like structure (**Supplementary Figs. 4d-f**). Inadequate Matrigel amounts also caused improper organoid formation (**Supplementary Figs. 4g-i**). Matrigel titration revealed that a concentration of approximately 50-100 µg/mL (equivalent to 1300-2700 ng per 25 µL droplet) was ideal for single AORB formation. These organoids exhibited a cell-free Matrigel core encased in a spherical epithelial shell, with diameters ranging from 484 ± 3.7 to 738 ± 11.2 µm (**Fig. 1g**).

Extensive characterization confirmed that differentiated AORBs had stably apical-out polarity, physiologically relevant cell compositions, transcriptional profiles, and orientation with apical-faced beating cilia and mucus production. Apical-out orientation of secretory cells was confirmed with a functional assay involving IL13-induced goblet cell hyperplasia, where gel-like substances visible outside AORBs in bright-field images were confirmed to contain mucins with fluorescent lectin. Additionally, the upregulation of *CLCA1* and *ALOX15B* were consistent with previous *in vitro* studies on IL13-induced *MUC5AC* expression^5–7^, suggesting similar cellular effects despite the geometrically inverted morphology. GO analysis demonstrated that the top 5 upregulated biological processes involved signaling pathways for ECM organization, cell-cell adhesion, and epithelial tube morphogenesis, while the top 5 downregulated biological processes related to cellular transcription/translational machinery. An additional reduction in proliferative gene expressions supported evidence of a differentiated state of cells in AORBs.

Do et al. reported no antiviral activity of AT-511 in 2D or 3D primary ALI cultures, a difference they attributed to be different experimental techniques/systems^8^. Our results in 2D transformed/immortalized cultures also showed little-to-no antiviral activity of AT-511^8^. Being a double prodrug, AT-511 requires a series of enzymatic processes to be activated, including cleavage of the ProTide moiety (via CatA, CES1, HINT1) and deamination of the base (ADALP1)^9^. As is seen for remdesivir, variation in enzyme levels between different cells will impact the level of active triphosphate (NTP) formed and the potency of the compound. Supporting this, preliminary *in vitro* assays show significantly higher levels of AT-511 processing into its active NTP form in HBTEC-ALI (170.8 pmol/10^6^ cells) compared with Vero (0.89 pmol/10^6^ cells), Caco-2 (30.1 pmol/10^6^ cells), and Calu-3 cells (4.93 pmol/10^6^ cells). Differences in potency and processing between 2D transformed vs. primary 3D cultures further highlight the importance of these models in the drug development pipeline.

Supplementary Table 1| AORB seeding and differentiation media compositions. O and X represent inclusion or exclusion of the component in seeding or differentiation media, respectively.

| **Media component** | **Supplier** | **Catalogue number** | **Final concentration** | **Seeding** | **Differentiation** |
| --- | --- | --- | --- | --- | --- |
| Cells | - | - | 120,000 cells/mL | O | X |
| Methylcellulose | Sigma-Aldrich | 94378 | 0.24% | O | X |
| Matrigel | Corning | 356231 | 50-100 µg/mL | O | X |
| Hydrocortisone | Sigma-Aldrich | 07925 | 1x | X | O |
| DAPT | Tocris | 2634 | 2 µM | X | O |
| PneumaCult^TM^ Airway Organoid Seeding Supplement | STEMCELL Technologies | 05062 | 10% | O | X |
| PneumaCult^TM^ Airway Organoid Basal Medium | STEMCELL Technologies | 05061 | 1x | O | O |
| PneumaCult^TM^ Airway Organoid Differentiation Supplement | STEMCELL Technologies | 05063 | 10% | X | O |

Supplementary Table 2| Total level of active nucleoside triphosphate (AT-9010).

|  | Vero^a^ | Caco-2^a^ | Calu-3^a^ | hBTEC-ALI^a^ |
| --- | --- | --- | --- | --- |
| AT-9010 (pmol/10^6^ cells) | 0.89 ± 0.05 | 30.1 ± 2.41 | 4.93 ± 0.06 | 170.8 ± 62.51 |
| ^a^Cells were treated with 10 μM of AT-511 for 4 h at 37°C. | | | | |

Supplementary Table 3. Antiviral response by the three donors. The overall variability attributes to the female donor, with more consistent results from other two male donors.

|  | **68YO, M** | | **65YO, F** | | **29YO, M** | |
| --- | --- | --- | --- | --- | --- | --- |
|  | **EC_50_^a^ (µM)** | **EC_90_ ^a^ (µM)** | **EC_50_ ^a^ (µM)** | **EC_90_ ^a^ (µM)** | **EC_50_ ^a^ (µM)** | **EC_90_ ^a^ (µM)** |
| Remdesivir | 0.3 ± 0.2 | 1.3 ± 0.9 | 1.8 ± 2.3 | 7.3 ± 8.3 | 0.4 ± 0.08 | 1.1 ± 0.7 |
| AT-511 | 1.1 ± 1.5 | 2.5 ± 0.9 | 0.8 ± 0.9 | 2.3 ± 1.1 | 0.3 ± 0.03 | 1.2 ± 0.04 |
| Nirmatrelvir | 0.08 ± 0.07 | 1.0 ± 0.8 | 0.1 | 1.0 | 0.3 ± 0.1 | 1.7 ± 1.4 |
| a EC = 50% (EC50) or 90% (EC90) effective antiviral concentration. | | | | | | |
